# *MYCN* Amplification and *ATRX* Mutations are Incompatible in Neuroblastoma

**DOI:** 10.1101/379636

**Authors:** Maged Zeineldin, Sara Federico, Xiang Chen, Yiping Fan, Beisi Xu, Elizabeth Stewart, Xin Zhou, Jongrye Jeon, Lyra Griffiths, John Easton, Heather Mulder, Donald Yergeau, Yanling Liu, Jianrong Wu, Collin Van Ryn, Arlene Naranjo, Michael D. Hogarty, Marcin M. Kamiński, Marc Valentine, Shondra M. Pruett-Miller, Alberto Pappo, Jinghui Zhang, Michael R. Clay, Armita Bahrami, Seungjae Lee, Anang Shelat, Rani E. George, Elaine R. Mardis, Richard K. Wilson, James R. Downing, Michael A. Dyer, for the St. Jude Children’s Research Hospital – Washington University Pediatric Cancer Genome Project

## Abstract

Aggressive cancers often have activating mutations in growth-controlling oncogenes and inactivating mutations in tumor-suppressor genes. In neuroblastoma, amplification of the *MYCN* oncogene and inactivation of the *ATRX* tumor-suppressor gene correlate with high-risk disease and poor prognosis. Here we show that *ATRX* mutations and *MYCN* amplification are mutually exclusive across all ages and stages in neuroblastoma. Using human cell lines and mouse models, we found that elevated *MYCN* expression and *ATRX* mutations are incompatible. Elevated MYCN levels promote metabolic reprogramming, mitochondrial dysfunction, reactive-oxygen species generation, and DNA-replicative stress. The combination of replicative stress caused by defects in the ATRX–histone chaperone complex and that induced by MYCN-mediated metabolic reprogramming leads to synthetic lethality. Therefore, *ATRX* and *MYCN* represent an unusual example, where inactivation of a tumor-suppressor gene and activation of an oncogene are incompatible. This synthetic lethality may eventually be exploited to improve outcomes for patients with high-risk neuroblastoma.

## INTRODUCTION

Neuroblastoma is the most common extracranial solid tumor of childhood^1^. *MYCN* amplification and age at diagnosis are the two most powerful predictors of outcome, with survival rates 5 to 10 times higher in infants than in adolescents or young adults (AYAs)^1–4^. Although *MYCN* amplification is equally distributed across age groups, previous genomic analyses of stage 4 pediatric neuroblastoma samples identified the *ATRX* mutations in 44% of AYAs (>12 y), 17% of children (18 mo–12 y), and 0% of infants (<18 mo)^1,5^. The patients whose tumors had *ATRX* mutations were typically older than 5 y, had an indolent disease course, and poor overall survival (OS).

One important function of ATRX is recognition of guanine (G)-rich stretches of DNA and deposition of the H3.3 histone variant to prevent the formation of G-quadruplex (G4) structures, which can block DNA replication or transcription^6–8^. G-rich repeats are found at telomeres and centromeres; ATRX forms a complex with DAXX to deposit H3.3 in those regions to maintain their integrity. Patients with ATR-X syndrome have aberrant chromatin at telomeres and pericentromeric DNA^7,8^. In tumor cells lacking ATRX, H3.3 is not deposited at the telomeric G-rich regions, G4 structures form, and replication forks stall^6–8^. As a result, telomeres undergo homologous recombination through the MRE11–RAD50–NBS1 (MRN) complex, leading to alternative lengthening of telomeres (ALT)^9^.

The formation of G4 structures in other G-rich repetitive regions of the genome can cause replicative stress^10–12^ or block transcription^13^. Indeed, H3.3 is deposited at actively transcribed genes in addition to telomeres and pericentromeric DNA^13^. ATRX may also affect transcription by targeting the PRC2 complex to particular regions of the genome^14^. As a result, in ATRX-deficient cells, PRC2-mediated modification of H3 to H3K27me3 lacks specificity, and genes that are normally repressed by polycomb are deregulated^14^.

MYCN regulates diverse cellular processes during development and in cancer. For example, elevated MYCN can lead to increased glycolytic flux and glutaminolysis in a variety of cancers including neuroblastomas^15–17^. MYCN also regulates genes important for fatty acid metabolism and nucleotide biosynthesis to promote metabolic reprogramming associated with tumorigenesis^15^. MYCN-induced glutaminolysis in neuroblastoma elevates reactive-oxygen species (ROS) and DNA-replicative stress^18,19^. As a result, neuroblastoma cells exhibit increased sensitivity to pharmacological agents that induce oxidative stress^18,19^. Here we demonstrate that the DNA-replicative stress induced by *ATRX* mutations and *MYCN* amplification cause synthetic lethality in neuroblastoma. This is unusual because oncogene activation and tumor-suppressor inactivation often work in concert to promote tumorigenesis not cancer cell death.

## RESULTS

### *ATRX* and *MYCN* mutations in neuroblastoma

To complement previous neuroblastoma studies from the Therapeutically Applicable Research to Generate Effective Treatment (TARGET) initiative^20^ and the Pediatric Cancer Genome Project (PCGP)^5,21^, we obtained neuroblastoma samples from 473 patients (122 unpaired and 351 paired tumor/germ line) from the Children’s Oncology Group (COG) (Table 1). We identified single-nucleotide variations (SNVs), small indels, and other somatic mutations in the coding region of *ATRX* via custom capture and Illumina sequencing using probes spanning the entire *ATRX* locus and whole-exome sequencing (WES) of 828 germline and tumor samples. We also included *MYCN* in the capture probe set to determine its copy number for each patient. We identified 19 somatic *ATRX* mutations and 15 of those are internal deletions predicted to encode truncated proteins (Fig. 1A).

**Figure 1.**
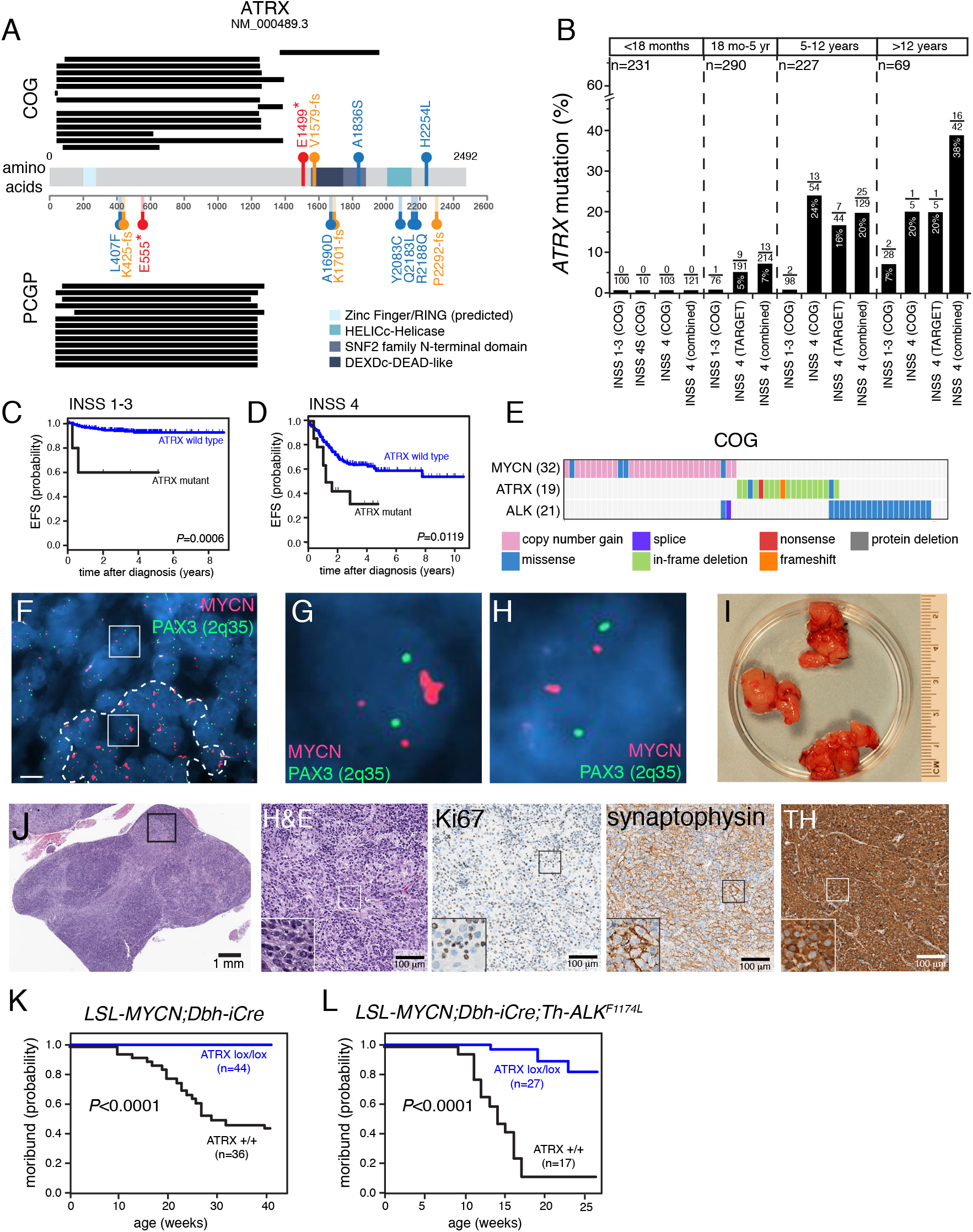
*ATRX* mutations are mutually exclusive of *MYCN* amplification in neuroblastoma. **A**) Summary of deletions that create ATRX^IFD^ proteins and singlenucleotide mutations in the COG and PCGP cohorts. Black bars indicate the deleted amino acids; red indicates nonsense mutations; orange indicates frameshift mutations; and blue indicates missense mutations. The major protein domains are shaded in blue. **B**) Bar plot of the percentage of the 819 patients analyzed in this study with *ATRX* mutations for each age and stage. The data are plotted from the COG, TARGET, and combined cohorts (PCGP cohort + COG cohort). **C,D**) EFS for patients with neuroblastoma with or without *ATRX* mutations for stages 1-3 or stage 4 for all age groups. **E**) Heatmap of the distribution of mutations in *MYCN, ATRX*, and *ALK* in the COG cohort. Only those tumors with a mutation in at least 1 of the 3 genes are indicated. **F-H**) Micrograph of *MYC* FISH (red) for the 1 neuroblastoma sample with an *ATRX* mutation and *MYCN* amplification showing regional amplification (dashed line). The PAX3 (2q35) probe (green) is the control. **G**) Micrograph of a neuroblastoma cell with *MYCN* homogenously staining region within a region of the tumor that has *MYCN* amplification (box within the dashed line region in **F**). **H**) Micrograph of a neuroblastoma cell with 2 copies of *MYCN* outside of the region with amplification (box outside of the dashed line region). **I**) Photograph of neuroblastoma tumors from a *LSL-MYCN;Dbh*-iCre mouse. **J**) Micrographs of the histology of the tumor shown in (I). The boxed region in each micrograph is magnified in the lower left panel of each image. **K,L**) Survival curves for 2 mouse models of neuroblastoma with or without conditional *ATRX* deletion. The numbers of mice in each group are indicated. Scale bar: F, 5 μm. **Abbreviations**: COG, Children’s Oncology Group; PCGP, Pediatric Cancer Genome Project; IDF, in-frame deletion; INSS, International Neuroblastoma Staging System; H&E, hematoxylin and eosin; TH, tyrosine hydroxylase.

Significantly higher ATRX-mutation frequencies were detected in patients with International Neuroblastoma Staging System (INSS) stage 4 disease (8.6%), high-risk subgroup (14.6%), 11q loss of heterozygosity, or unfavorable histology and in those who were 18 mo or older at diagnosis (Table 1, Fig. 1B). *ATRX*-mutant neuroblastomas were significantly more likely to have ALT than were *ATRX* wild-type tumors [89.5% (17/19) vs. 22.2% (4/18; p <0.0001, 2-sided Fisher’s exact test]. Additionally, black patients had a significantly higher frequency of *ATRX* mutation than did those in other racial categories (9.1% vs. 3.3%; p=0.0348) (Table 1). *ATRX* mutation frequency did not differ based on sex, *ALK* status, tumor grade, mitosis-karyorrhexis index, or ploidy (Table 1).

*ATRX* mutations were associated with significantly lower 4-year event-free survival (EFS) rates among the groups with non–INSS stage 4 disease (p=0.0007), INSS stage 4 disease (p=0.0128), ages 5–12 y (p=0.0006) and older than 12 y (p=0.0038) at diagnosis in the COG cohort (Table S1 and Fig. 1C,D). Compared to those without a mutation, patients harboring an *ATRX* mutation were 5.0 times more likely to experience an adverse event (p<0.0001) and 3.4 times more likely to die of their disease (p=0.0046) (Table S1).

In the cohort of 819 tumors, *ATRX* mutations (n=64) and *MYCN* amplification (n=140) showed statistically significant mutual exclusivity (p=0.0037, Cochrane-Mantel-Haenszel test) (Fig. 1E and Supplemental Information). Only 1 of 819 (0.1%) tumors had both lesions. However, this was an unusual discordant sample because it was scored as MYCN-amplified by fluorescence in situ hybridization (FISH) in COG’s neuroblastoma reference lab but nonamplified based on normalized read depth from our custom-capture Illumina next-generation sequence (NGS) data (Fig. S1A). To resolve this discrepancy, we repeated the FISH analysis and found regional heterogeneity of *MYCN* amplification (Fig. 1F-H). Specifically, the region with *MYCN* amplification had a high mitosis-karyorrhexis index and was less likely to have ultrabright telomere foci, which are characteristic of ALT in *ATRX*-mutant neuroblastoma cells (Figs. 1F-H and S1B-E).

To determine if *ATRX* mutations and *MYCN* amplification are incompatible in vivo, we conditionally inactivated *ATRX* in 2 genetically engineered mouse models of neuroblastoma (Fig. 1I-L)^11,22,23^. One model (*LSL-MYCN;Dbh-iCre*) is a transgenic line with conditional expression of *MYCN* from the *Rosa26a* locus mediated by Cre expressed in the dopaminergic cells of the sympathoadrenal lineage^22^. The other model includes the *Th-ALK^F1174L^* transgene that potentiates MYCN-mediated tumorigenesis^23^. For each of these models, tumor formation was reduced and survival was significantly increased (p <0.0001) when *ATRX* was simultaneously inactivated with elevated *MYCN* expression (Fig. 1K-L).

### *ATRX* mutations are incompatible with elevated *MYCN*

To extend our data from mice to humans, we characterized the expression of MYCN, ATRX, and DAXX proteins across 12 human cancer cell lines (Fig. 2A). We used 8 neuroblastoma cell lines, including 4 lines with *MYCN* amplification (IMR-32, SKNBE2, NB-5) or moderately elevated levels of MYCN (NBL-S) (Table S2), 2 lines (SKNMM and CHLA90) produce ATRX with an in-frame deletion (ATRX^IFD^) and are diploid for *MYCN* (Fig. 2A and data not shown), one osteosarcoma cell line with no detectable expression of ATRX (U2OS) and 3 lines as controls (293T, HeLa, and SJSA1)^24^.

**Figure 2.**
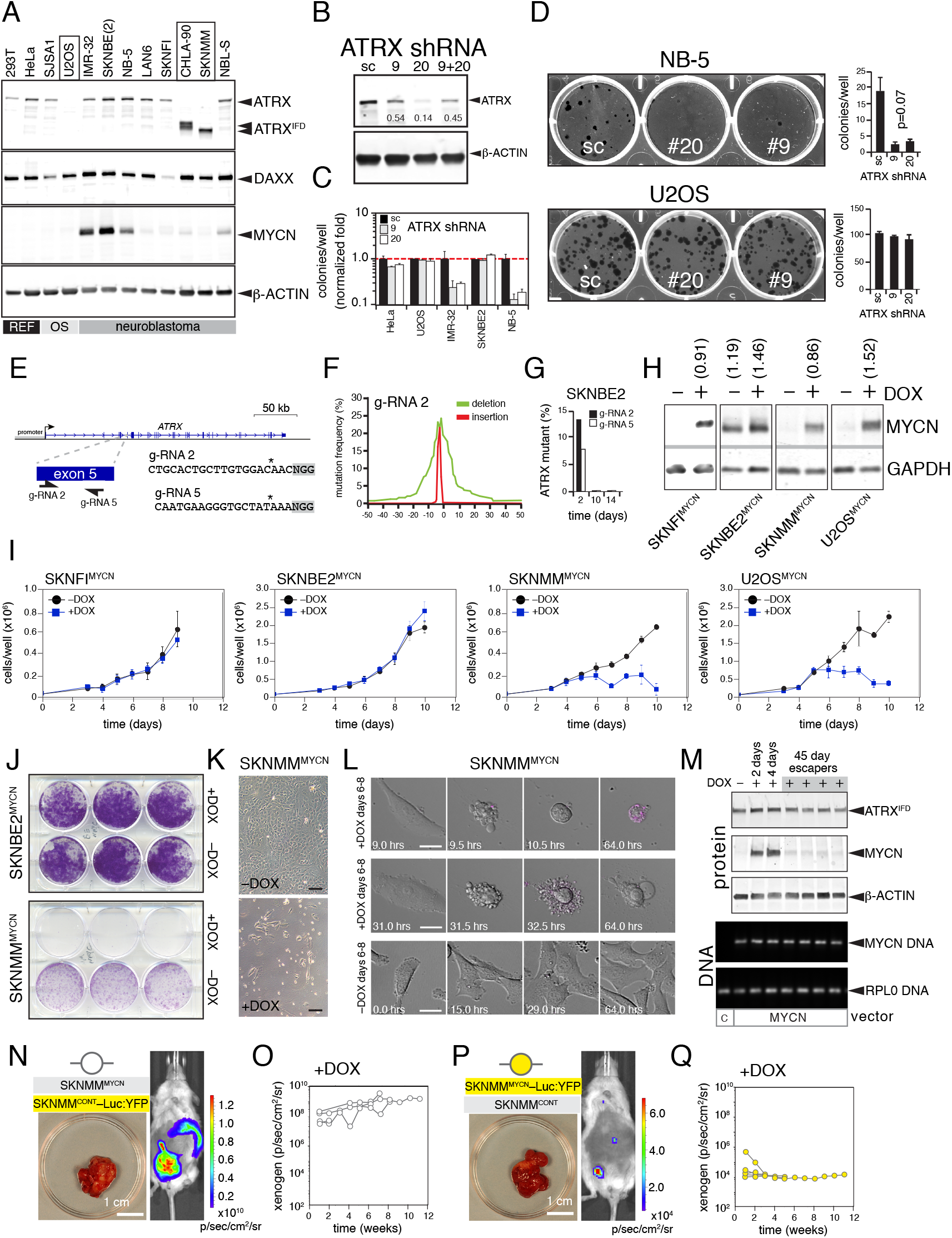
*ATRX* mutations and *MYCN* amplification are incompatible. **A**) Immunoblot of ATRX, DAXX, MYCN and β-actin across 12 cell lines. Boxes indicate ATRX mutant lines. **B**) Immunoblot for ATRX and β-actin in 293T cells transfected with sc shRNA or ATRX-specific shRNAs (#9, #20). The fraction of ATRX remaining relative to sc shRNA is indicated on each lane. **C**) Bar plot of the number quantity of colonies per well after transfection of each shRNA as normalized relative fold to the sc shRNA (dashed line). Each bar is the mean and standard deviation of duplicate experiments. **D**) Photograph of cresyl violet–stained colonies of NB5 and U2OS cells after transfection with the shRNAs and bar plots of the number of colonies per well for duplicate experiments (mean and standard deviation). **E**) Map of 2 gRNAs targeting exon 6 of *ATRX*. **F**) Plot of the frequency of mutation for gRNA 2, with deletions in green and insertions in red, from MiSeq analysis of the PCR product spanning the target sequence of *ATRX*. **G**) Bar plot of the proportion of MiSeq reads from the PCR product spanning the target sequence for gRNA 2 and gRNA 5 in SKNBE2 cells after 2, 10, or 14 days in culture. **H**) Immunoblot for MYCN and GAPDH for the cell lines with stable integration of the doxycycline-inducible MYCN-expression construct. Numbers in parentheses above the lanes indicate the fluorescence-intensity ratio of MYCN/GAPDH. **I**) Line graphs of the growth curves for each cell line in the presence (blue) or absence (black) of doxycycline. **J**) Colony assay in the presence or absence of doxycycline for SKNBE2^MYCN^ and SKNMM^MYCN^ cells. **K**) Brightfield micrograph of SKNMM^MYCN^ cells after 8 days in culture in the presence or absence of doxycycline. **L**) Brightfield micrographs of 3 individual cells or a field of cells for SKNMM^MYCN^ in the presence or absence of doxycycline at the indicated timepoints, relative to the starting point of the movies at 6 days after the addition of doxycycline. **M**) Immunoblots of ATRX, MYCN, and β-actin in SKNMM^MYCN^ cells without doxycycline or after 2, 4, or 45 days in culture. The escapers are pools of cells that grew in the presence of doxycycline. PCR for the MYCN transgene and a control locus (RPL0) is indicated in the lower portion of the figure. **N**) Xenogen image of a mouse with an orthotopic neuroblastoma tumor, which is shown in the photograph, that arose after para-adrenal injection of a 1:1 mixture of SKNMM^MYCN^ cells and SKNMM^CONT^–Luc:YFP cells. **O**) Line graph of xenogen image data for individual mice after the injection described in (**N**). **P**) Xenogen image of a mouse with an orthotopic neuroblastoma tumor, which is shown in the photograph, that arose after para-adrenal injection of a 1:1 mixture of SKNMM^MYCN^–Luc:YFP cells and SKNMM^CONT^ cells. **Q**) Line graph of xenogen image data for individual mice after the injection described in (**P**). Scale bars: K, 10 μm. **Abbreviations**: DOX, doxycycline; IFD, inframe deletion; sc, scrambled.

To knock down ATRX protein, we used 2 different lentiviral vectors expressing shRNAs targeting *ATRX* (#9 and #20) (Fig. 2B). The optimal knockdown was approximately 86% in HeLa cells after 48 h with ATRX shRNA–20; similar results were achieved across the other 11 cell lines (data not shown). Two weeks after introducing shRNAs, we measured telomere fluorescence per cell and performed telomere Q-PCR for each cell line. Neither telomere content nor ALT status differed after ATRX knockdown in any cell line tested (Fig. S2A-H and data not shown). The proportions of cells in G0/G1, S, or G2/M phases of the cell cycle did not differ across all cell lines tested (Fig. S2I). Knocking down ATRX decreased colony formation in MYCN-amplified neuroblastoma cells (IMR-32 and NB-5) but not cell lines expressing wild-type *MYCN* (Figs. 2C,D).

To inactivate ATRX in the MYCN-amplified neuroblastoma cell lines, we designed and validated 2 guide RNAs (gRNA-2 and gRNA-5) to exon 6 of *ATRX* (Fig. 2E,F). We transfected 293T, SKNBE2, and IMR32 cells with each gRNA and a plasmid expressing Cas9 and purified the transfected cells by flow cytometry. At 2, 10, and 14 days after plating the purified cells, we harvested DNA, performed PCR using primers spanning both of the 2 gRNA-target sequences, and performed Illumina MiSeq analysis to calculate the proportion of mutated alleles in each sample at each timepoint. Although the *ATRX*-mutant alleles were abundant in 293T and SKNBE2 cells 2 days after plating, those cells harboring the mutated alleles of *ATRX* were lost by 10 days in culture (Figs. 2G and S2J). In MYCN-amplified IMR32 cells, no mutant alleles were detected at any timepoint, though a control gRNA to the *AAVS1* locus led to efficient mutagenesis (Fig. S2J). A similar study performed with a gRNA to *DAXX* had no effect on the viability of 293T, SKNBE2, or IMR32 cells (Fig. S2K-M and data not shown).

Although these results suggested that reducing the expression of *ATRX* is incompatible with *MYCN* amplification in neuroblastoma, we could not isolate and study the cells because they died. Therefore, we induced the expression of *MYCN* in *ATRX-* mutant neuroblastoma cells by using a doxycycline-inducible vector (Fig. 2H and supplemental information). Initially, we used 2 *ATRX*-mutant cell lines (SKNMM and U2OS), a MYCN-amplified cell line (SKNBE2), and a cell line that is wild-type for both genes (SKNFI) (Fig. 2A). The levels of MYCN achieved with doxycycline induction in both ATRX-mutant lines were similar to those in MYCN-amplified cell lines (Fig. 2H).

The growth rates of SKNFI^MYCN^ and SKNBE2^MYCN^ cells, after 10 days in culture ± doxycycline, showed no difference with induction of MYCN expression, but by 6 days in culture, the 2 ATRX-mutant lines showed a dramatic loss of cells (Fig. 2I). A parallel experiment performed with stable cell lines that had only the doxycycline inducible vector lacking *MYCN* showed no effect on cell growth (data not shown). A similar result was observed in colony assays (Fig. 2J and data not shown). More than 95% of the cells in the SKNMM^MYCN^ cultures were lost by 8–10 days in the presence of doxycycline (Fig. 2K). Live-imaging of cells ± doxycycline on Days 6–8, the peak period of cell loss (Fig. 2L; movies available upon request), enabled us to determine the timing of cell death, relative to cytokinesis, in 44 cells from 13 different movies (Table S3). In total, 54% of the cells died after initiating cytokinesis; the remaining 46% had a long period (20.2 ± 12.8 h) between cytokinesis and cell death (Table S3). These data were extended to 3 other ATRX-mutant cell lines (Fig. S3A,B). There was no effect of ectopic MYCN expression on a DAXX mutant cell line (Fig. S3A,B).

A small number of SKNMM^MYCN^ cells escaped cell death and continued to grow for 45 days in the presence of doxycycline. The ATRX^IFD^ was still expressed and the *MYCN* transgene was still present, but MYCN protein was lost (Fig. 2M).

To determine if an elevated level of MYCN is incompatible with *ATRX* mutation in vivo, we performed an in vivo growth competition assay. In 1 cohort of mice, SKNMM^MYCN^ cells labeled with luciferase and YFP were mixed in a 1:1 ratio with unlabeled control SKNMM^CONT^ cells, which contained the doxycycline-inducible vector lacking *MYCN*. In a second cohort, the SKNMM^CONT^ cells were labeled with luciferase and YFP and the SKNMM^MYCN^ cells were unlabeled. Cells were injected into the paraadrenal space of 25 immunocompromised (NOD-SCID) mice for each cohort. The luciferase was monitored every week, and ultrasound was performed every 2 weeks. Animals were randomized to +doxycycline and –doxycycline groups once the tumor was larger than 14 mm^3^ by 3-dimensional ultrasound and/or 1.4×10^7^ photons/s/cm^2^/sr by xenogen imaging (Table S4). The SKNMM^MYCN^ cells were outcompeted by the SKNMM^CONT^ cells in vivo, as determined by xenogen, ultrasound, and flow cytometry (Fig. 2N-Q and data not shown).

### *MYCN* targets in neuroblastoma

MYCN is a global transcriptional regulator that activates and represses genes that control cell division, cell size, and cell differentiation during development^25^. MYCN can upregulate gene expression by heterodimerizing with MAX and binding the CACGTG E-box sequence at the promoters of its target genes^25^. In addition, MYCN can up- or downregulate genes by recruiting histone-modifying or chromatin-remodeling complexes^25,26^. To analyze the epigenomic landscape in *ATRX*-mutant neuroblastoma cells, compared to that in wild-type *ATRX* neuroblastoma, we selected 8 cell lines that encompass the major genetic groups, including those with *MYCN* amplification or *ATRX* mutations (Fig. 3A). We also included 8 orthotopic patient-derived xenografts (O-PDXs) that were derived by para-adrenal injection of patient tumors into immunocompromised mice as described previously^27,28^ (Fig. 3A). One of those O-PDXs (SJNBL047443_X1) has an *ATRX* in frame deletion (Fig. 3B,C and Table S2). Autopsy tumor material from a patient with ATRX-mutant neuroblastoma (SJNBL030014_D) was also obtained for transcriptome, epigenomic, and genomic profiling (Fig. 3A and Table S2). We performed whole-genome bisulfite sequencing (WGBS; Fig. S4 and Table S5), RNA-sequencing, and ChIP-seq of 8 histone marks (H3K4me1, H3k4me2, H3K4me3, H3K27me3, H3K27Ac, H3K36me3, H3K9/14Ac, and H3K9me3), CTCF, BRD4, and RNA polymerase II (PolII) across all 8 cell lines, the 8 O-PDX tumors, the autopsy sample, and normal fetal adrenal medulla. We also performed MYCN ChIP-seq on 3 O-PDXs with *MYCN* amplification (SJNBL046_X, SJNBL012407_X1, and SJNBL013762_X1) and the SKNFI^MYCN^, SKNBE2^MYCN^, and SKNMM^MYCN^ cell lines with doxycycline on Day 4 in culture as well as matched controls. To relate gene expression changes to epigenetic changes induced by MYCN, we performed ChIP-seq for the 8 histone marks, CTCF, Brd4, and RNA PolII in the SKNMM^MYCN^ cells ± doxycycline on Day 4 in culture as described above.

To define the chromatin states and analyze the transitions thereof across the genome, we performed chromatin Hidden Markov Modeling (ChromHMM) using all 393 ChIP-seq datasets (Fig. 3D,E and Table S6). To share these data, we have developed a cloud-based viewer (https://pecan.stjude.org/proteinpaint/study/mycn_nbl_2018) that displays the gene expression, ChromHMM, WGBS, and ChIP-seq tracks for each sample. We identified 394 genes that were upregulated and 544 that were downregulated 4 days after adding doxycycline to the SKNMM^MYCN^ or U2OS^MYCN^ cell lines (Table S7). Gene-set-enrichment analysis of the upregulated genes identified MYC and MYCN targets as the most enriched datasets (Fig. S5A): 84% of the upregulated genes and 42% of the downregulated genes had a MYCN-binding site based on MYCN ChIP-seq for SKNMM^MYCN^ cells in the presence of doxycycline (Fig. S5B-E and Table S7). Analysis of the chromHMM for the upregulated MYCN target genes showed that expansion of state 13 (strong transcribed gene body) downstream of the transcriptional start site was the most significant change (Supplemental Information, Fig. 3F,G and Table S7). This is consistent with previous data showing that MYC regulates RNA PolII pause release not transcriptional initiation^29^. These upregulated target genes are enriched for the pathways involved in metabolism (e.g., *SLC3A2, SLC7A5*) and mitochondrial gene expression (e.g., *PNPT1*) (Figs. 3 F,G, S5B-E, and Table S7), which is consistent with previous studies demonstrating that elevated MYC/MYCN induces metabolic reprogramming in cancer cells^15^.

**Figure 3.**
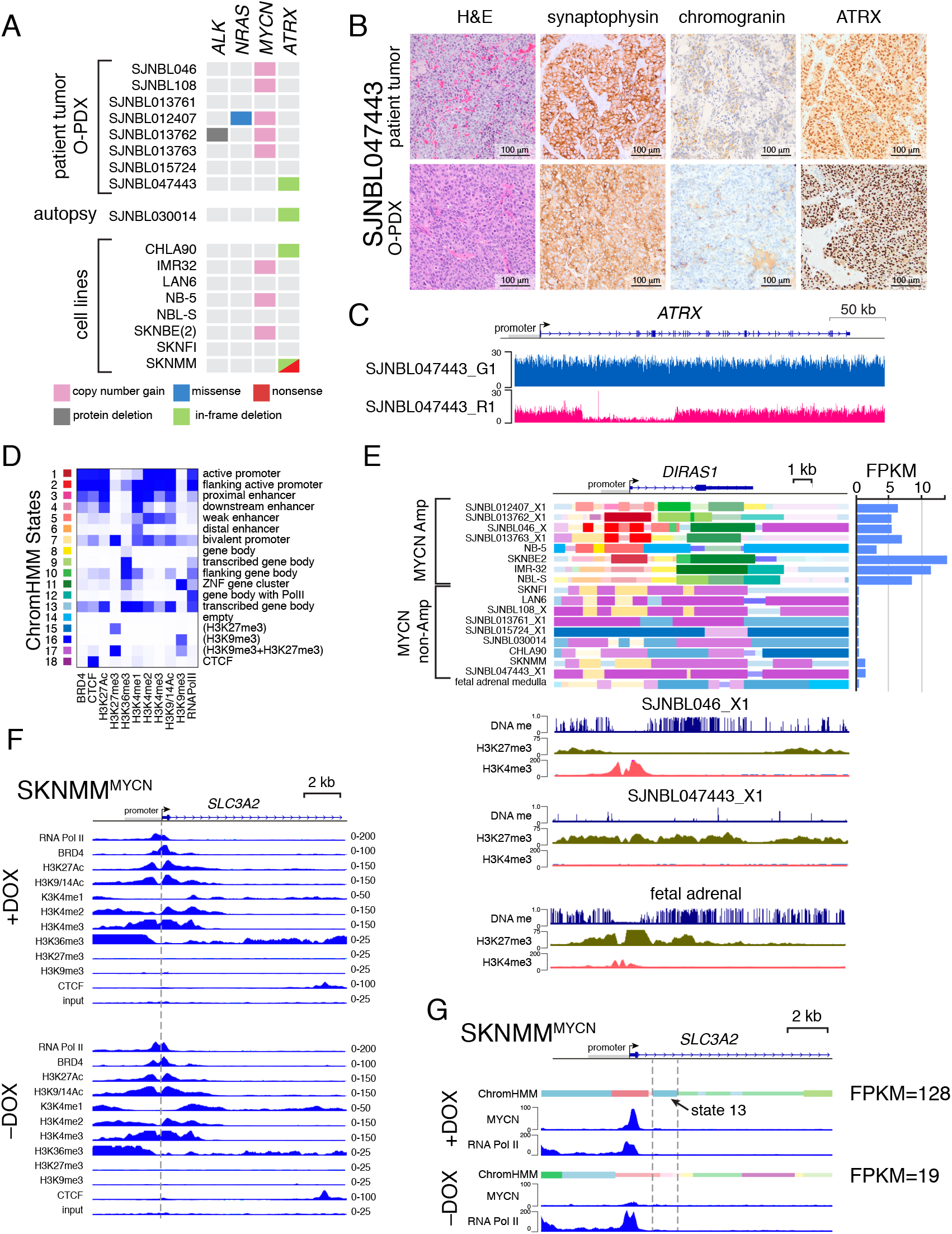
Transcriptional and epigenetic changes induced by MYCN in *ATRX-* deficient neuroblastoma cells. **A**) Heatmap of the mutations in the O-PDXs and cell lines used in this study. An autopsy sample was also used for molecular profiling. **B**) Micrographs of histologic analysis of the *ATRX*-mutant neuroblastoma O-PDX and corresponding patient tumor. **C**) Plot of sequence read depth from Illumina sequencing for the *ATRX* gene in the patient’s germline (blue) and tumor (red), showing somatic deletion of exons 2-11. **D**) Heatmap of the chromHMM states used in this study with functional annotation. **E**) ChromHMM plot of the *DIRAS1* gene and a bar plot of the gene’s expression on the right. DNA methylation and H3K27me3 and H3K4me3 ChIP-seq for *DIRAS1* for a MYCN-amplified tumor (SJNBL046_X1), an ATRX-mutant tumor (SJNBL047443_X1), and fetal adrenal medulla. **F**) ChIP-seq tracks for a portion of the *SLC3A2* gene spanning the promoter in SKNMM^MYCN^ cells in the presence or absence of doxycycline after 4 days in culture. Dashed line indicates the start of transcription. The scales are indicated on the right. **G**) ChromHMM and ChIP-seq for MYCN and RNA PolII for a portion of the *SLC3A2* gene spanning the promoter in SKNMM^MYCN^ cells in the presence or absence of doxycycline after 4 days. The dashed lines indicate the region with induction of ChromHMM state 13 that is expanded after MYCN binds the promoter. **Abbreviations**: ChromHMM, chromatin Hidden Markov Modeling; FPKM, fragments per kilobase of transcript per million mapped reads; O-PDX, orthotopic patient-derived xenograft.

### Metabolic reprogramming induced by MYCN

One form of metabolic reprogramming induced by MYC/MYCN is glutamine addiction^30^. The glutamine transporter (SLC1A5) involved in glutamine uptake and the bidirectional transporter (SLC7A5–SLC3A2) involved in glutamine efflux coupled with essential amino acid import are direct targets of MYCN and are upregulated in SKNMM^MYCN^ cells in the presence of doxycycline (Table S7, Figs. 3G and S5D,E)^31^. Following *MYCN* induction, we quantified glutamine metabolism in SKNMM^MYCN^ cells ± doxycycline on Day 4 in culture and measured 8 tricarboxylic acid (TCA) cycle cellular metabolites^32^. The proportions of 5-carbon–labeled glutamate and a-ketoglutarate increased, consistent with glutamine utilization for the TCA cycle as well as reductive carboxylation to produce 5-carbon labeled citrate (Fig. 4A,B).

**Figure 4.**
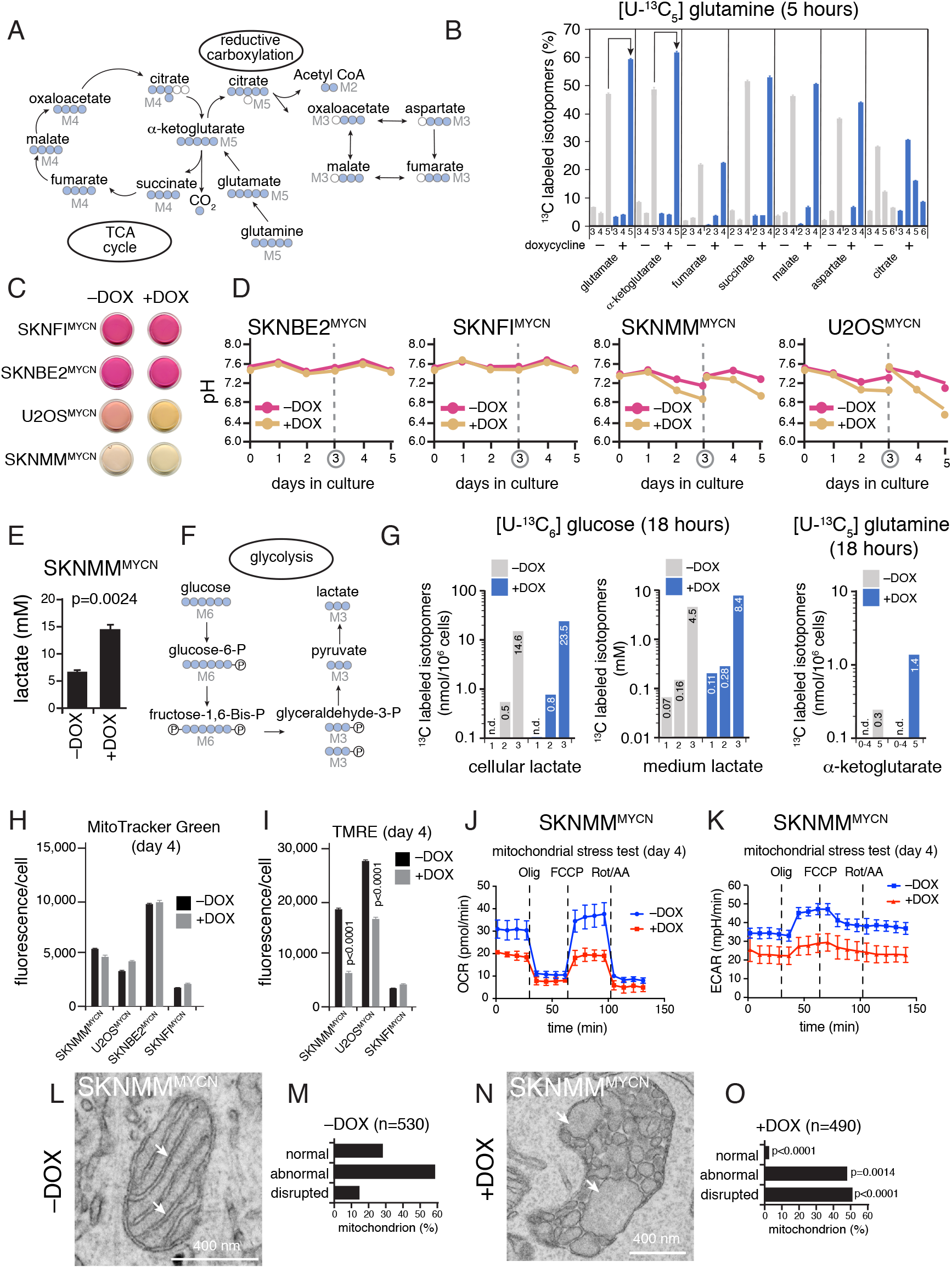
Expression of MYCN in ATRX-mutant cells leads to metabolic reprogramming and mitochondrial dysfunction. **A**) Simplified drawing of the TCA cycle and reductive carboxylation to highlight the metabolites that were measured. The blue circles indicate the radiolabeled carbons derived from uniformly labeled ^13^C_5_-glutamine by each pathway. **B**) Bar plot of ^13^C-labeled isotopomers in SKNMM^MYCN^ cells after 4 days in the presence or absence of doxycycline and 5 hours of labeling with uniformly labeled ^13^C_5_-glutamine. The arrows indicate an increase in glutamate and α-ketoglutarate in the presence of doxycycline. Each bar is the mean and standard error of the mean of technical replicates. **C**) Photograph of culture media 3 days after the addition of doxycycline for each cell line. SKNFI and SKNBE2 cells are maintained in culture in Dulbecco’s modified eagle medium; U2OS cells are maintained in McCoy’s medium; and SKNMM cells are maintained in RPMI. D) Line plots of pH for each of the 4 cell lines in the presence or absence of doxycycline for each day in culture. The culture media were changed on Day 3. **E**) Histogram of the levels of lactate at Day 4 in SKNMM^MYCN^ cells with or without doxycycline. Each bar is the mean and standard deviation of biological replicate experiments. **F**) Simplified drawing of the glycolysis pathway highlighting the production of lactate from uniformly labeled ^13^C_6_–glucose. **G**) Bar plots of ^13^C-labeled isotopomers of lactate in the medium or cells after 18 h of labeling with uniformly labeled ^13^C_6_–glucose at Day 4 in SKNMM^MYCN^ cells with or without doxycycline. The concentrations are indicated on the bars. Similar data are shown for α-ketoglutarate after 18 h of labeling with uniformly labeled ^13^C_5_–glutamine. **H,I**) Bar plots of MitoTracker Green staining (**H**) and TMRE (**I**) staining as the geometric mean and 95% confidence interval of the fluorescence per cell in the presence or absence of doxycycline on Day 4 in culture. **J,K**) Representative measurements of oxygen consumption rate (OCR) and extracellular acidification rate (ECAR) with 2.5×10^4^ cells grown under the Cell Mito Stress assay conditions for the seahorse. **L-O**) Representative electron micrograph of mitochondria from SKNMM^MYCN^ cells after 4 days in culture in the presence or absence of doxycycline (**L,N**) showing disruption in the mitochondrial cristae (arrows). Bar plot of scoring of mitochondrial morphology for the SKNMM^MYCN^ cells in the presence or absence of doxycycline after 4 days in culture (**M,O**). **Abbreviations**: DOX, doxycycline; FCCP, carbonyl cyanide-p-trifluoromethoxy phenylhydrazone; Olig, oligomycin; Rot/AA, rotenone/antimycin A; TCA, tricarboxylic acid; TMRE, tetramethylrhodamine ethyl ester.

Inefficient use of glutamine and/or glucose by cancer cells can lead to the export of glycolytic intermediates such as lactate^33^. The pH of the medium was quickly acidified in subconfluent cultures of SKNMM^MYCN^ or U2OS^MYCN^ cells in the presence of doxycycline; however, SKNBE2^MYCN^ and SKNFI^MYCN^ cells showed no difference in the pH of their media (Fig. 4C,D). The reduced pH correlated with an increase in lactate in the culture medium and dependence on glutamine for survival (Fig. 4E and Fig. S6A). To determine whether the lactate was produced from glutamine or glucose and provide additional data on the metabolic reprogramming induced by MYCN in ATRX-deficient cells, we performed a more comprehensive metabolomic profiling of 54 metabolites using ^13^C-labeled glucose and ^13^C-labeled glutamine (Fig. 4F,G, Supplemental Information, and Table S8). Glucose was the major source of lactate; relatively few of the carbon from glucose was used for the TCA cycle (Figs. 4F,G and Table S8). In contrast, glutamine was converted to a-ketoglutarate for the TCA cycle and reductive carboxylation (Figs. 4F,G and Table S8).

In addition to its role in nucleotide and amino acid synthesis, glutamine is an important mitochondrial substrate; cells must precisely balance the expression and activity of proteins encoded by the nucleus with those encoded by the mitochondria to maintain homeostasis^32^. Cells also upregulate pathways required to mitigate the reactive oxygen species (ROS) that are a natural biproduct of mitochondrial metabolism to prevent excessive protein or DNA damage. For example, glutamine uptake in cancer cells can elevate glutathione levels because glutamine is converted to glutamate, which is a precursor for glutathione synthetase^34^. We measured the levels of glutathione for each cell line ± doxycycline; cell lines that were sensitive to glutamine depletion in the medium also had significantly elevated levels of glutathione (p <0.0001, Fig. S6B). ROS and DNA damage were increased in the absence of glutamine (Fig. S6C,D).

Pathways involved in mitochondrial homeostasis were upregulated in SKNMM^MYCN^ or U2OS^MYCN^ cells in the presence of doxycycline (Table S7). Little change in mitochondrial mass was measured using MitoTracker Green in SKNMM^MYCN^ or U2OS^MYCN^ cells ± doxycycline, but the mitochondrial membrane potential (Δψ_m_) measured with tetramethylrhodamine ethyl ester (TMRE) was significantly lower in the presence of doxycycline (p <0.0001; Fig. 4H,I). To assess mitochondrial function in MYCN-overexpressing SKNMM cells, we measured the oxygen consumption rate. Induction of MYCN in SKNMM cells decreased basal mitochondrial respiration and respiratory reserve capacity, relative to that seen in uninduced SKNMM cells (Figs. 4J,K and S7). The scoring of mitochondrial ultrastructure on transmission electron micrographs showed that SKNMM^MYCN^ cells in the presence of doxycycline had more disrupted mitochondria on Day 4 and subsequent timepoints than did SKNMM^MYCN^ cells maintained in culture without doxycycline or the other cell lines (Figs. 4L-O, S8 and data not shown).

### Replicative stress in ATRX-mutant neuroblastomas expressing MYCN

The mitochondrial dysfunction described above may lead to the accumulation of ROS and in turn contribute to replicative stress through DNA damage. We measured ROS and DNA damage in the SKNMM^MYCN^, SKNBE2^MYCN^, SKNFI^MYCN^, and U2OS^MYCN^ cells ± doxycycline. The ROS increase began at Day 4 and persisted until Day 6 in SKNMM^MYCN^ cells in the presence of doxycycline (Fig. 5A and data not shown). This was accompanied by increased expression of γH2AX protein, a biomarker for DNA double-strand breaks (Fig. 5B,C). Spectral karyotyping showed a significant increase in cells with DNA fragmentation and translocation in the SKNMM^MYCN^ cells in the presence of doxycycline (p <0.0001; Fig. 5D). We also detected increased DNA fragmentation in SKNMM^MYCN^ and U2OS^MYCN^ cells in the presence of doxycycline by using a single-cell gel electrophoresis assay (Fig. 5E-G and data not shown).

**Figure 5.**
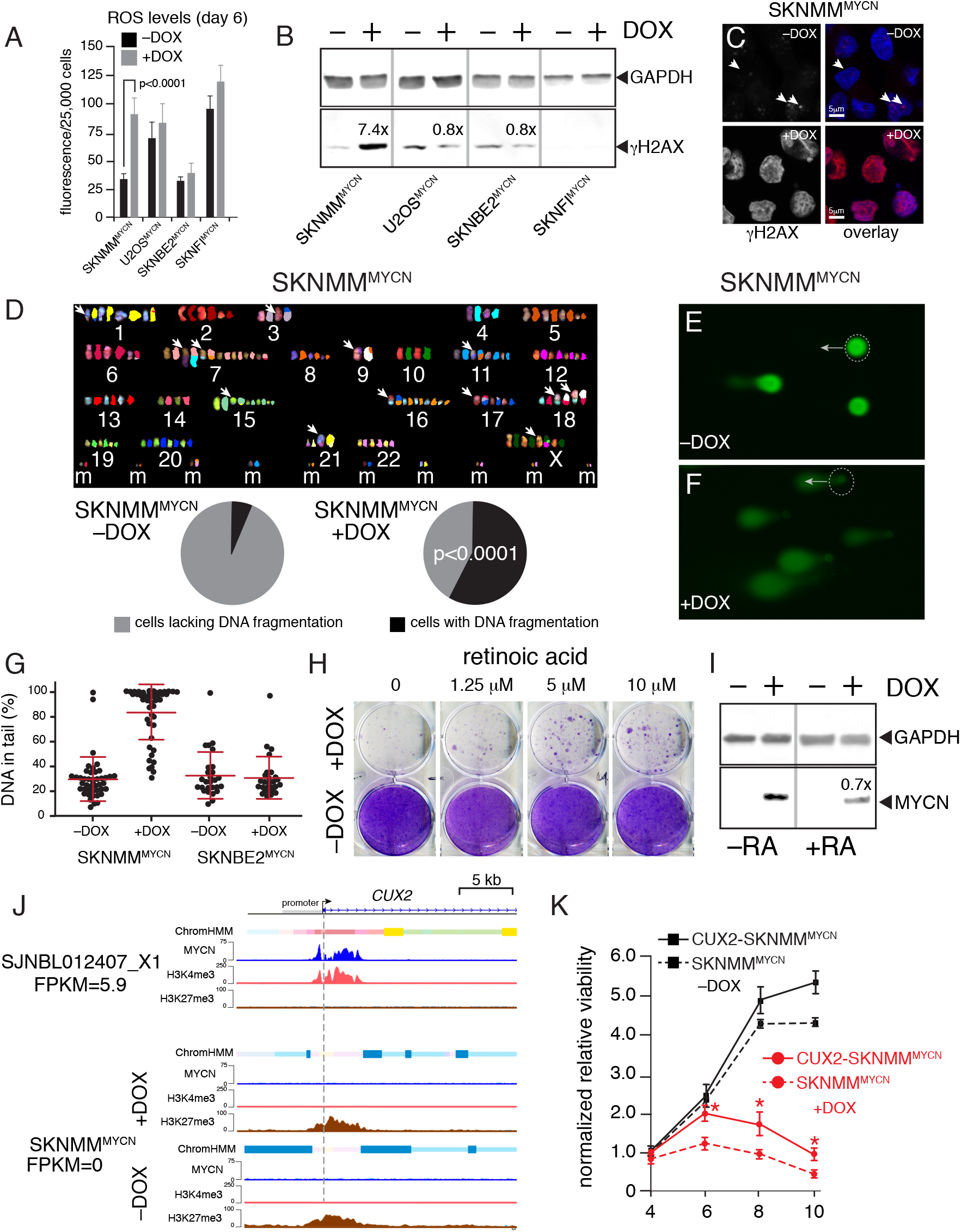
Expression of MYCN in ATRX-mutant cells leads to reactive-oxygen species production, DNA damage, and replicative stress. **A**) Bar plot of ROS levels on Day 6 in culture for 4 cell lines in the presence or absence of doxycycline. Each bar is the mean and standard deviation of triplicate samples. **B**) Immunoblots of γH2AX and GAPDH on Day 4 in the presence or absence of doxycycline. **C**) Immunofluorescent detection of γH2AX (red) and nuclei (blue) in SKNMM^MYCN^ cells on Day 4 in culture in the presence or absence of doxycycline. **D**) Spectral karyotype analysis (SKY) of SKNMM^MYCN^ cells on Day 8 in culture. Each chromosome is shown adjacent to the pseudo-colored representation. Arrows indicate chromosomes with translocations. The pie charts show the proportion of cells with DNA fragmentation, as detected using SKY. E,F) Micrographs of single-cell electrophoresis of individual nuclei (dashed circles) and their COMET tail (arrows) of SKNMM^MYCN^ cells in the presence or absence of doxycycline on Day 5 in culture. **G**) Scoring of the COMET assay shown in (**E,F**). **H**) Photograph of cresyl violet–stained colonies from SKNMM^MYCN^ cells in the presence or absence of doxycycline, with increasing concentrations of retinoic acid (RA). **I**) Immunoblots of GAPDH and MYCN in SKNMM^MYCN^ cells in the presence or absence of doxycycline, with or without 5 μM RA. The level of MYCN protein was reduced by 30% (0.7x) in the presence of RA. **J**) ChromHMM and ChIP-seq for MYCN, H3K4me3, and H3K27me3 for the *CUX2* promoter for a MYCN-amplified neuroblastoma (SJNBL012407_X1) xenograft and SKNMM^MYCN^ cells in the presence or absence of doxycycline after 4 days in culture. The gene expression (FPKM) is indicated, and the dashed line indicates the start of transcription. **K**) Line plot of SKNMM^MYCN^ cells in the presence or absence of doxycycline after 4, 6, 8 and 10 days in culture with or without ectopic expression of *CUX2* or the GFP control from lentiviral infection. Each point is the mean and standard deviation of triplicate experiments, and the asterisks indicate statistical significance (p<0.05). **Abbreviations**: DOX, doxycycline; RA, retinoic acid; ROS, reactive-oxygen species.

Cells with DNA replicative stress are sensitive to hydroxyurea^35^. Thus, we maintained SKNMM^MYCN^ cells cultured ± doxycycline with different concentrations of hydroxyurea (12.5, 25, 50, or 100 μM on Days 1–4) and measured cell viability using CellTiter-Glo on Day 4. The SKNMM cells expressing MYCN were significantly more sensitive to hydroxyurea at 50 or 100 μM concentrations (p<0.0001; Fig. S9A). The DNA content in each cell line, ± doxycycline at Day 4 in culture, showed an increase in S and G2/M phases in the SKNMM^MYCN^ cells in the presence of doxycycline, which was consistent with cell cycle arrest as a result of DNA damage (Fig. S9B). The ROS scavenger N-acetyl cysteine had a small but reproducible effect on reducing cell death induced by ectopic MYCN expression in SKNMM cells (Fig. S9C,D).

To identify other pathways that may modulate to MYCN-induced cell death in ATRX-deficient cells, we performed a dose-response screen using 2 drug libraries. The first was a collection of 177 oncology drugs and compounds in late clinical development (Phase II or later). The second was a customized library of 34 drugs and compounds that target proteins or pathways that have been reported to reduce MYC/MYCN oncogenic activity (Table S9). Within the oncology drug set, 37 compounds selectively potentiated killing of SKNMM^MYCN^ cells in the presence of doxycycline (Table S9); the majority (59.4%, 22/37) were drugs that induce DNA damage, inhibit DNA repair or replication, or inhibit cell cycle checkpoints (Fig. S9E-G and Table S9). The only 2 drugs that partially rescued the cell death were a third-generation retinoid derivative (bexarotene) and an mTORC1/2 inhibitor (INK128) (Fig. S9H and Table S9). The bexarotene result was particularly interesting because retinoids are an effective treatment for neuroblastoma (i.e., they reduce MYCN expression and induce differentiation). These results were independently validated using retinoic acid (Fig. 5H,I).

Together, these data suggest that MYCN induction causes metabolic reprogramming and mitochondrial dysfunction that contribute to increased ROS and DNA-replicative stress. It may be synthetically lethal in *ATRX*-mutant cells because they already have DNA-replicative stress in part due to reduced ability to resolve G4 DNA structures. A subset of genes that are normally induced in MYCN-amplified neuroblastomas also may have failed to be activated in the *ATRX*-mutant neuroblastomas due to the DNA and/or histone modifications. We identified 1 gene that met those criteria. *CUX2* encodes a homeodomain protein with 3 CUT repeats that is expressed in the developing nervous system and important for the repair of oxidative DNA damage^36^. *CUX2* is a direct MYCN target and is upregulated in MYCN-amplified neuroblastomas but not ATRX-mutant neuroblastomas (Fig. 5J). The *CUX2* promoter is epigenetically repressed by H3K27me3 in *ATRX*-mutant neuroblastomas and not induced in SKNMM cells in the presence of doxycycline (Fig. 5J). To determine if CUX2 is important for oxidative stress in MYCN-amplified neuroblastomas, we ectopically expressed the protein in SKNMM^MYCN^ cells ± doxycycline. There was a partial rescue of cell death in SKNMM^MYCN^ cells in the presence of doxycycline when CUX2 was ectopically expressed (Fig. 5K).

## DISCUSSION

Mutually exclusive mutation profiles are not uncommon in cancer cells; however, many are thought to result from targeting the same oncogenic pathway^37^. For example, inactivation of the *RB1* tumor-suppressor gene is often exclusive of amplification of genes encoding cyclins (e.g., *CCNE1*) because they target the same pathway. Examples of synthetic lethality caused by oncogene activation and tumor-suppressor mutation are much less common, and few (if any) have been validated in vivo.

Here we show that amplification of the *MYCN* oncogene and inactivation of the *ATRX* tumor-suppressor gene are mutually exclusive in neuroblastomas from patients of all ages and stages of disease. One small, discrepant tumor sample may have contained 2 separate clones, but detailed analysis was not possible. In mouse models and cell lines, the combination of elevated MYCN expression and ATRX loss led to synthetic lethality. Ectopic MYCN caused metabolic reprogramming, mitochondrial dysfunction, ROS production, and DNA damage in ATRX-mutant neuroblastoma cells. We propose that *MYCN* amplification and *ATRX* mutations are incompatible in neuroblastoma, because both lead to DNA-replicative stress^10,38,39^. Consistent with this model, the synthetic lethality was partially rescued by genes that reduce oxidative stress (*CUX2*) and pharmacological agents that induce differentiation (retinoic acid) or reduce ROS levels (N-acetyl cysteine). Based on data presented here, this synthetic lethality may extend to other MYC-driven tumors.

*ATRX*-mutant neuroblastomas have several unique features, relative to other ATRX-mutant cancers. First, *DAXX* was not mutated in our cohort, whereas in pancreatic neuroendocrine tumors (PanNETs), *ATRX* and *DAXX* mutations have approximately the same frequency and are mutually exclusive^38^. Second, *ATRX*-mutant neuroblastomas have worse outcome, but patients with ATRX/DAXX-mutant PanNETs tumors have prolonged survival^38^. Third, *ATRX* mutations in neuroblastomas are in-frame deletions that remove approximately half of the amino terminus of the protein. In other cancers, such as PanNETs, the mutations are indels or nonsense mutations^38^.

Previous studies suggest that amino acids 1-841 of ATRX are sufficient for localization to heterochromatin^39^. This region is deleted in the ATRX^IFD^s in most *ATRX*-mutant neuroblastomas. The ADD domain (amino acids 168-293) binds to histone H3 tails, with preferential binding to H3K9me3 domains that lack H3K4 methylation. HP1 can bind H3K9me3 and recruit ATRX through the HP1-interacting domain (amino acids 586-590), even in the absence of the ADD domain in ATRX. Therefore, in neuroblastomas with in-frame deletions, we propose that they lack the heterochromatin-binding domain and have defects in H3.3-chaperone function that is required for genome maintenance. We cannot rule out other mechanisms that may contribute to the synthetic lethality, such as modulation of gene expression through PRC2^14^ or resolution of G4 DNA structures for DNA replication or transcription.

Some tumor cell lines with *ATRX* mutations are hypersensitive to ATR inhibitors^40^. This may be exploited for the treatment of patients with neuroblastoma whose tumors have *ATRX* mutations. However, the more promising and challenging approach is to develop strategies to reproduce the cell-intrinsic reprogramming that was achieved with MYCN overexpression and *ATRX* deletion in neuroblastoma cells in our studies. If successful, this could potentially benefit patients with MYCN-amplified tumors by reducing the function of ATRX. Alternatively, inducing metabolic (or other changes) caused by MYCN expression in ATRX-mutant tumors could be useful. To achieve this ambitious goal of generating synthetic lethality in neuroblastoma cells in patients with high-risk disease by targeting these 2 pathways, much will need to be learned about the downstream targets of ATRX and MYCN that contribute to this phenotype.

## ACKNOWLEDGEMENTS

We thank Angie McArthur for editing the manuscript and Kevin Freeman for the MYCN-inducible constructs. The NBL-S cell line was provided by Garrett Brodeur. This work was supported, in part, by Cancer Center Support (CA21765) from the NCI, grants to M.A.D from the NIH (EY014867, EY018599, and CA168875) and ALSAC. M.A.D. was also supported by a grant from Alex’s Lemonade Stand Foundation for Childhood Cancer, the Tully Family Foundation, and the Peterson Foundation. E.S. was supported by the St. Baldrick’s Foundation. The statistical work was supported by NIH/NCI grant U10 CA180899 (Children’s Oncology Group Statistics and Data Center). Most of this research was supported by the Howard Hughes Medical Institute. WGS, WGBS, and RNA-seq were performed as part of the St. Jude Children’s Research Hospital – Washington University PCGP. Preclinical studies were performed with assistance from the Animal Imaging Shared Resource; electron microscopy was performed with assistance from the Cell and Tissue Imaging Shared Resource; histopathologic analysis was performed with assistance from the Veterinary Pathology Shared Resource (all of St. Jude).

## METHODS

### Statistical Analysis of *ATRX* Mutations in Neuroblastoma

Patient samples were grouped according to the INSS for stage and age at diagnosis: infants, toddlers, children, and AYA. The frequency of *ATRX* mutations was calculated across the cohort, in subgroups based on age, and INSS categories (stage 1, 2A/B or 3, or 4S and 4)^2^. Chi-square tests or Fisher’s exact tests were used determine associations between *ATRX* mutations and clinical variables. EFS and OS were compared between patients with or without *ATRX* mutations by using a log-rank test. EFS and OS Cox models were used to assess the prognostic strength of *ATRX* mutations in the presence of standard risk factors. EFS was measured as the number of days from diagnosis to date of relapse, disease progression, secondary malignancy, death, or, if no event occurred, date of last contact. OS was measured as the number of days from diagnosis to date of death, or if the patient had not died, date of last contact.

### Animals and O-PDX Tumors

All animal procedures and protocols were approved by the St. Jude Laboratory Animal Care and Use Committee. NOD/SCID/gamma and CD1-nude mice were purchased from the Jackson Laboratory. Human O-PDXs were generated and expanded as described previously^41^. All O-PDX tumors described here are freely available with no obligation to collaborate through the Childhood Solid Tumor Network (http://www.stjude.org/CSTN/).

### Sequence Data Access

All sequence data were deposited in the EMBL-EBI database.

Additional materials and methods are available in the Supplemental Information.

---

**Figure S1.**
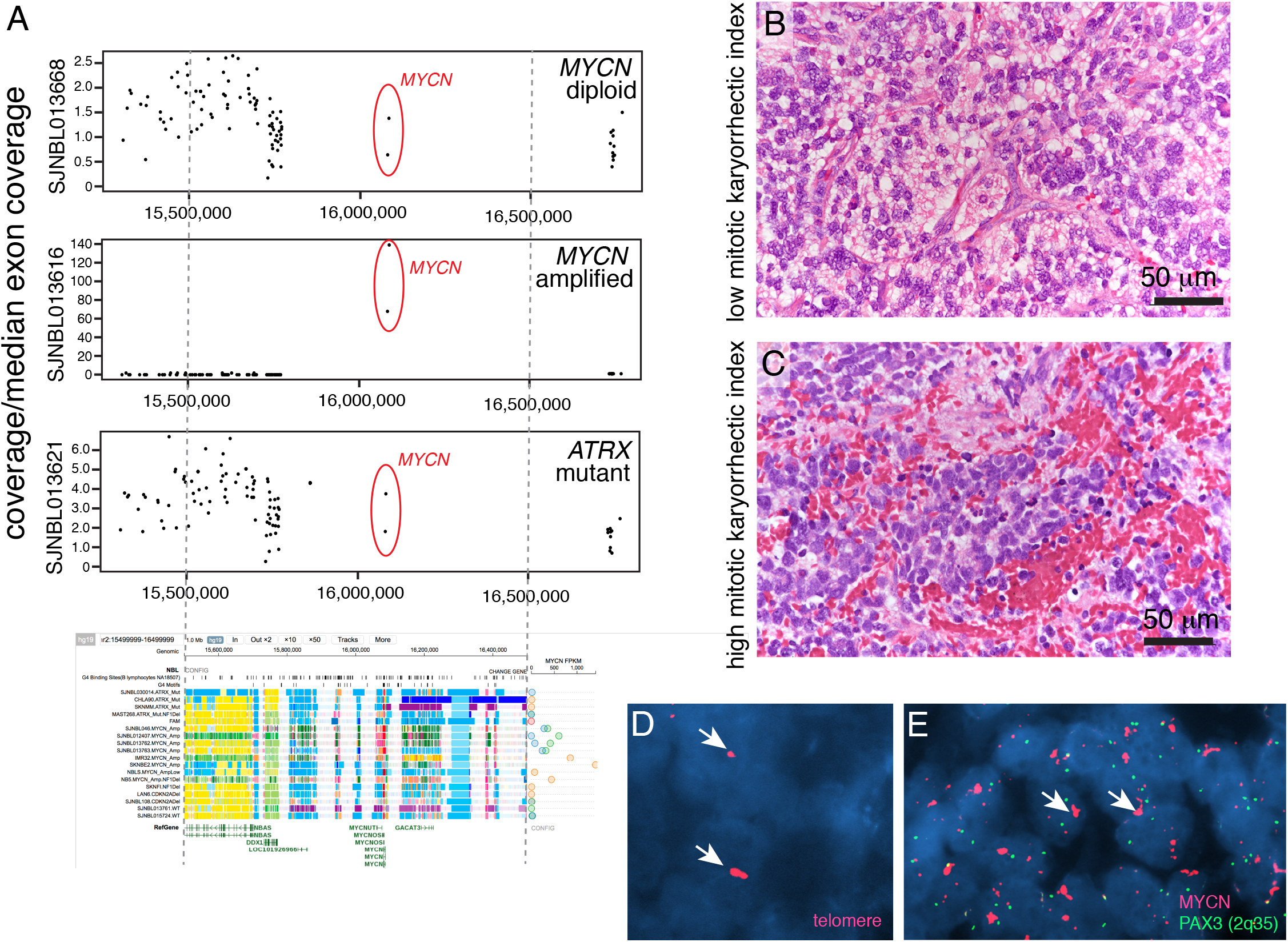
Analysis of *MYCN* amplified *ATRX* mutant sample. **A**) Scatterplot of the coverage relative to the median exon coverage for the 3 indicated samples. SJNBL013668 has the normal diploid copy number of the MYCN locus, SJNBL013616 has high level amplification and SJNBL013621 is the sample reported to have MYCN amplification and *ATRX* mutation. The genomic region is aligned with the ChromHMM viewer that is part of this study showing the location of the exons being plotted. **B,C**) Micrograph of Hematoxylin and Eosin staining of the SJNBL013621 sample showing regions of low and high mitotic karyorrhectic index. **D**) Micrograph of fluorescent in situ hybridization (FISH) for telomeres (red) with nuclear stain (blue) showing regions of ultrabright signal indicative of alternative lengthening of telomeres (ALT). **E**) Micrograph of FISH for *MYCN* (red) and a control probe (*PAX3*, green). Arrows indicate strong FISH signals for telomeres and *MYCN*.

**Figure S2.**
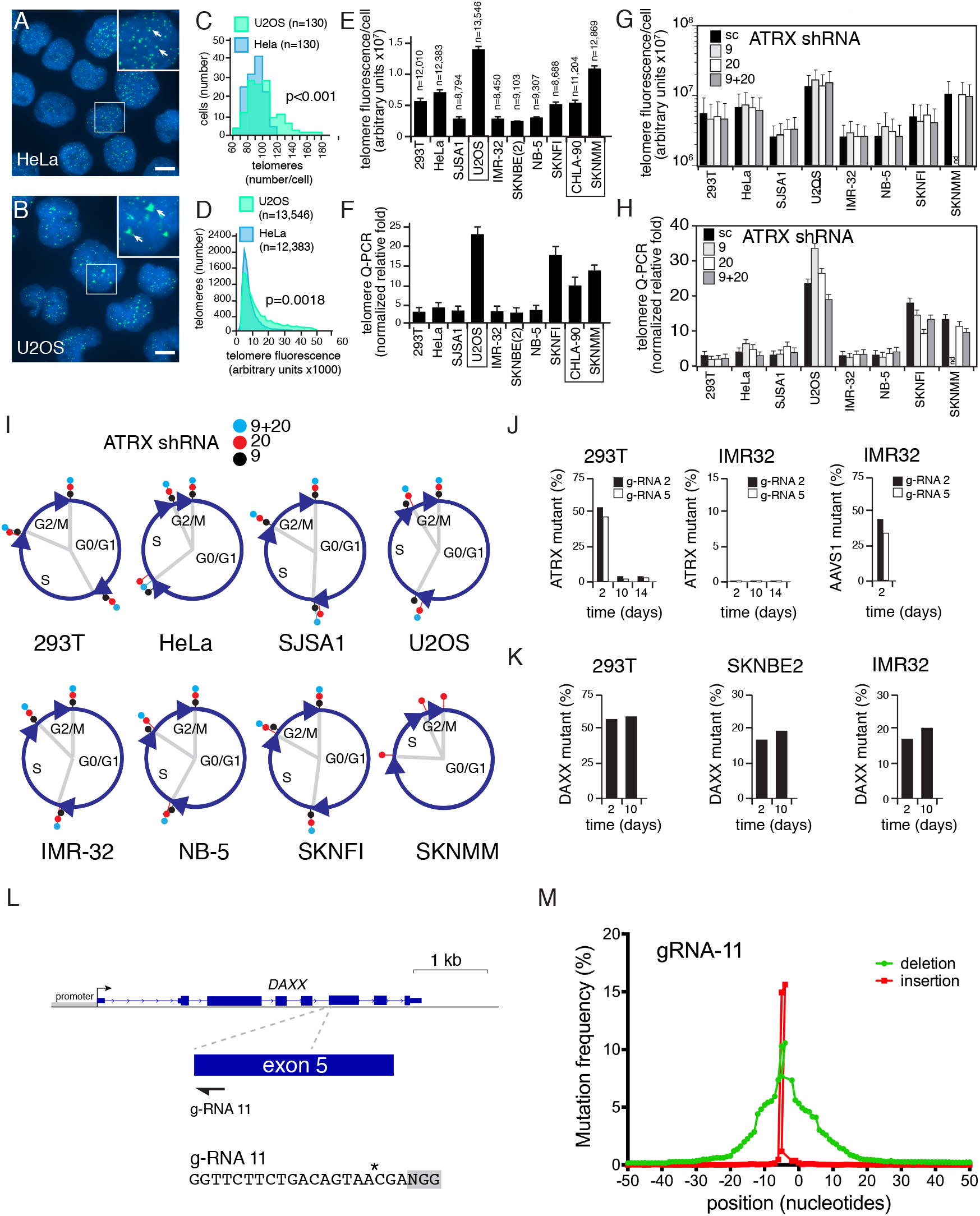
Inactivation of ATRX and DAXX. **A,B**) Micrograph of telomere FISH for HeLa and U2OS cells showing individual telomeres (arrows). **C**) Step plot of the number of telomeres per cell for HeLa cells (blue) and U2OS cells (green). **D**) Distribution of fluorescence intensity of individual telomeres in HeLa and U2OS cells. The ultrabright telomere signal in the U2OS (arrows in panel B) and higher intensity per telomere (green in D) are hallmarks of alternative lengthening of telomeres (ALT). **E**) Bar plot of telomere fluorescence per cell for the indicated number of telomeres for each cell line **F**) Bar plot of the amount of telomere DNA as measured by QPCR for each of the cell lines. **I**) Piecharts of the cell cycle distribution for each of the shRNAs for each cell line. **J**) Proportion of MiSeq reads from the PCR product spanning the target sequence for g-RNA 2 and g-RNA 5 in 293T, SKNBE2 and IMR32 cells after 2, 10 and 14 days in culture. A control for IMR32 (AAVS1) is shown to demonstrate that efficient CRISPR/Cas9 targeting can be achieved in those cells. **K**) Proportion of MiSeq reads from the PCR product spanning the target sequence for g-RNA11 targeting exon 5 of DAXX. **L**) Map of the location of g-RNA 11 targeting exon 5 of DAXX. **M**) Plot of the frequency of mutation for g-RNA 2 with deletions in green and insertions in red from MiSeq analysis of the PCR product spanning the target sequence of DAXX. Scale bar: 5 μm.

**Figure S3.**
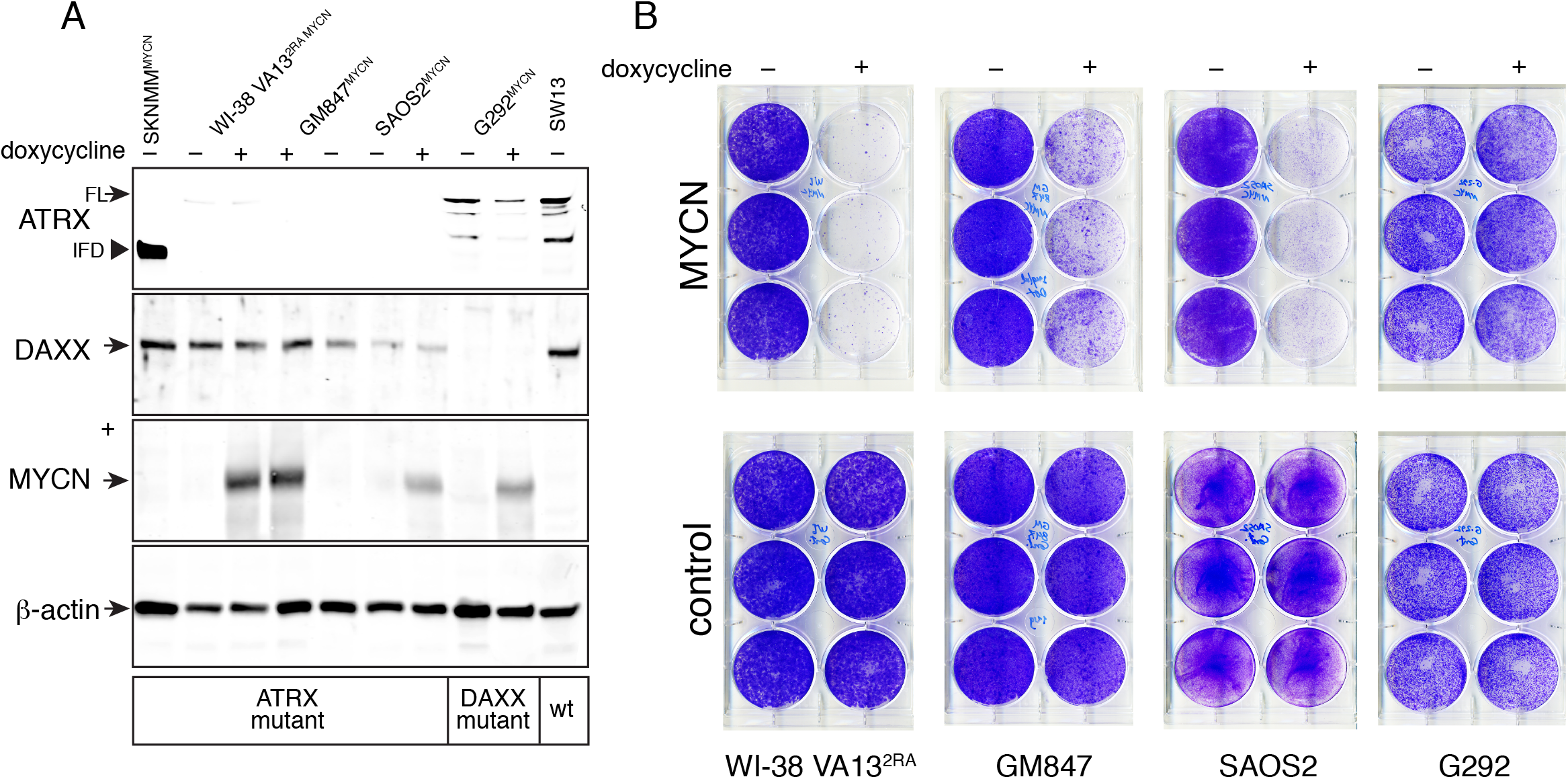
Analysis of additional ATRX and DAXX deficient cell lines. **A**) Immunoblot of 6 cell lines that have stable integration of the doxycycline inducible MYCN expression plasmid. Lysates were prepared 4 days after addition of doxycycline and blotted for ATRX, DAXX, MYCN and β-actin as a loading control. SKNMM cells have an in frame deletion of ATRX (IFD), WI-38 VA13^2RA^, GM847 and SAOS2 cells lack ATRX while G292 cells lack DAXX. The SW13 cells were included as a wild type control for ATRX and DAXX. **B**) Photograph of a cresyl violet stained 6 well dish from colony assays with each of the indicated cell lines in the presence and absence of doxycycline. The control lines have the tet regulated plasmid integrated into the genome but lacking the MYCN transgene.

**Figure S4.**
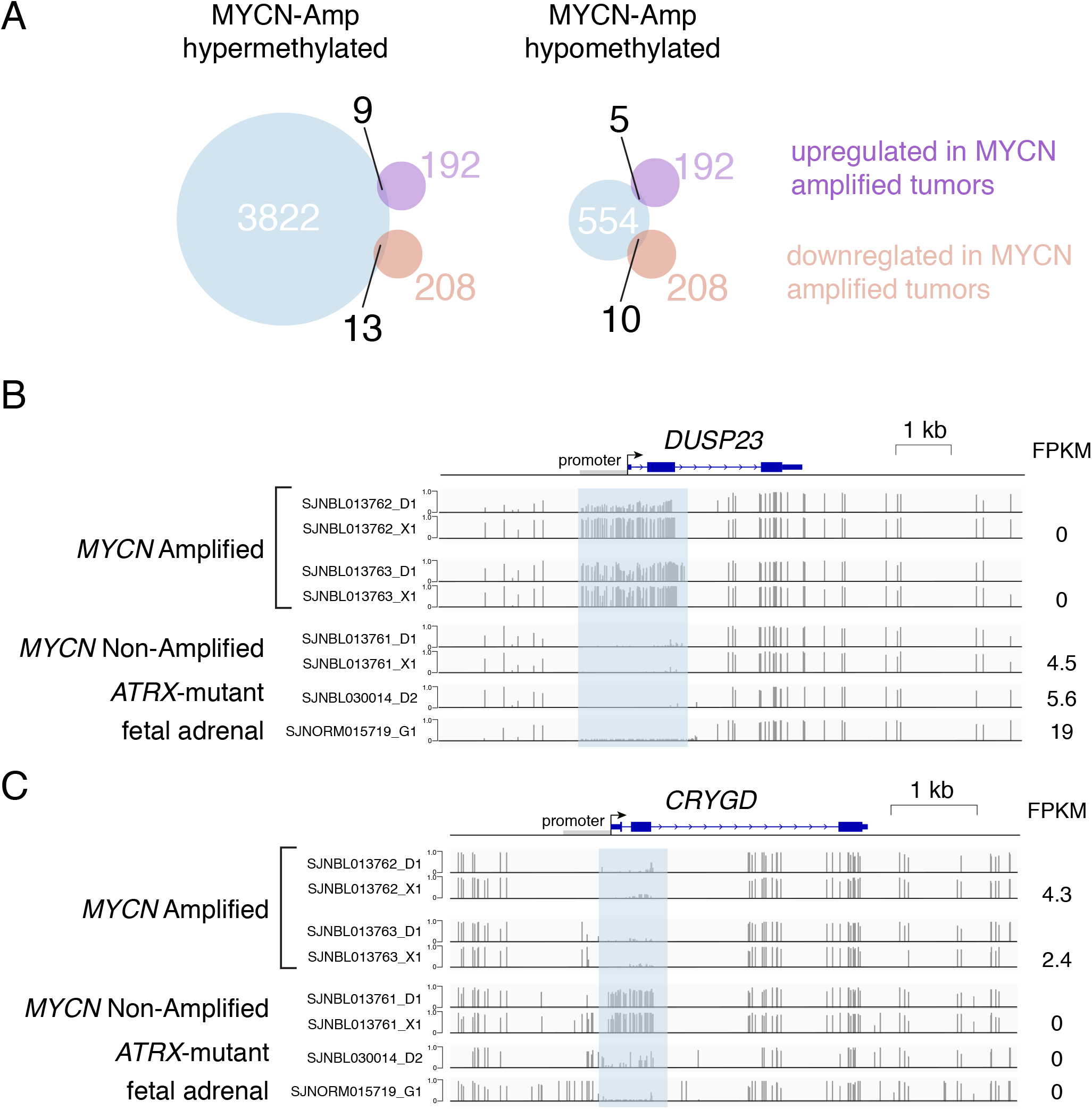
DNA methylation analysis. Venn diagram of the hypermethylated and hypomethylated promoters (blue) in the *MYCN* amplified tumors relative to the non-amplified tumors. The upregulated genes are shown in purple and the downregulated genes are shown in brown. The number of overlapping genes are indicated on the diagram. **H,I**) Representative plots of DNA methylation from whole genome bisulfite sequencing for a representative hypermethylated promoter (*DUSP23*) in *MYCN* amplified tumors that correlated with reduced gene expression (FPKM is shown to the right) and a hypomethylated promoter (*CRYGD*) that correlated with elevated gene expression. Abbreviations: FPKM, fragments per kilobase of transcript per million reads.

**Figure S5.**
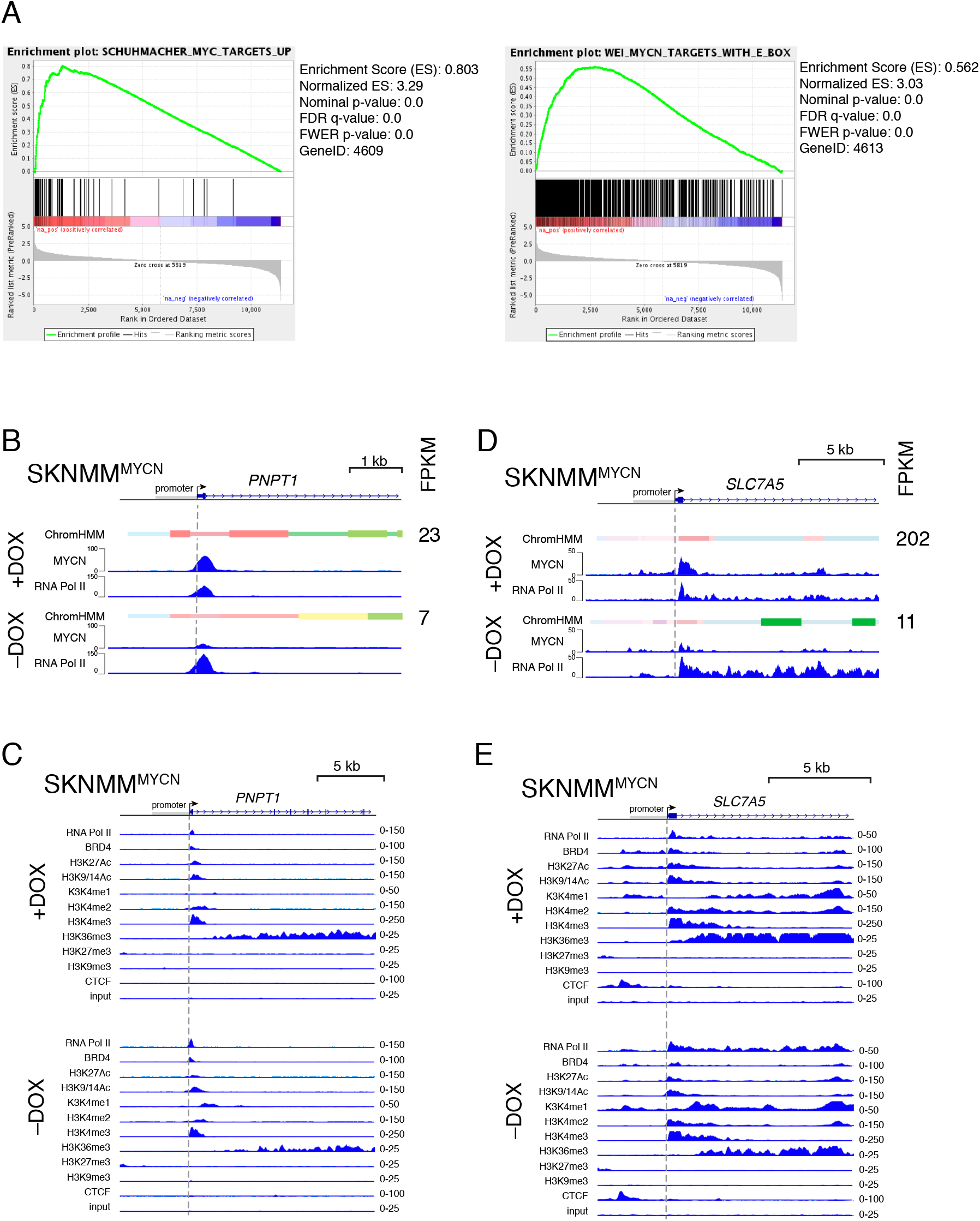
Induction of MYCN targets in SKNMM^MYCN^ cells. **A**) The top pathways in gene set enrichment analysis for SKNMM^MYCN^ and U2OS^MYCN^ cells in the presence of doxycycline relative to the absence of doxycycline. **B)** ChIP-seq for MYCN and RNA Pol II and chrom-HMM for the *PNPT1* gene promoter in SKNMM^MYCN^ cells in the presence and absence of doxycycline. The PNPT1 protein is important for importing nuclear encoded RNA into the mitochondria. **C**) All ChIP-seq tracks for the *PNPT1* gene in the presence and absence of doxycycline. **D**) ChIP-seq for MYCN and RNA Pol II and chrom-HMM for the *SLC7A5* gene promoter in SKNMM^MYCN^ cells in the presence and absence of doxycycline. The SLC7A5 protein is important for glutamine transport. **E**) All ChIP-seq tracks for the *SLC7A5* gene in the presence and absence of doxycycline.

**Figure S6.**
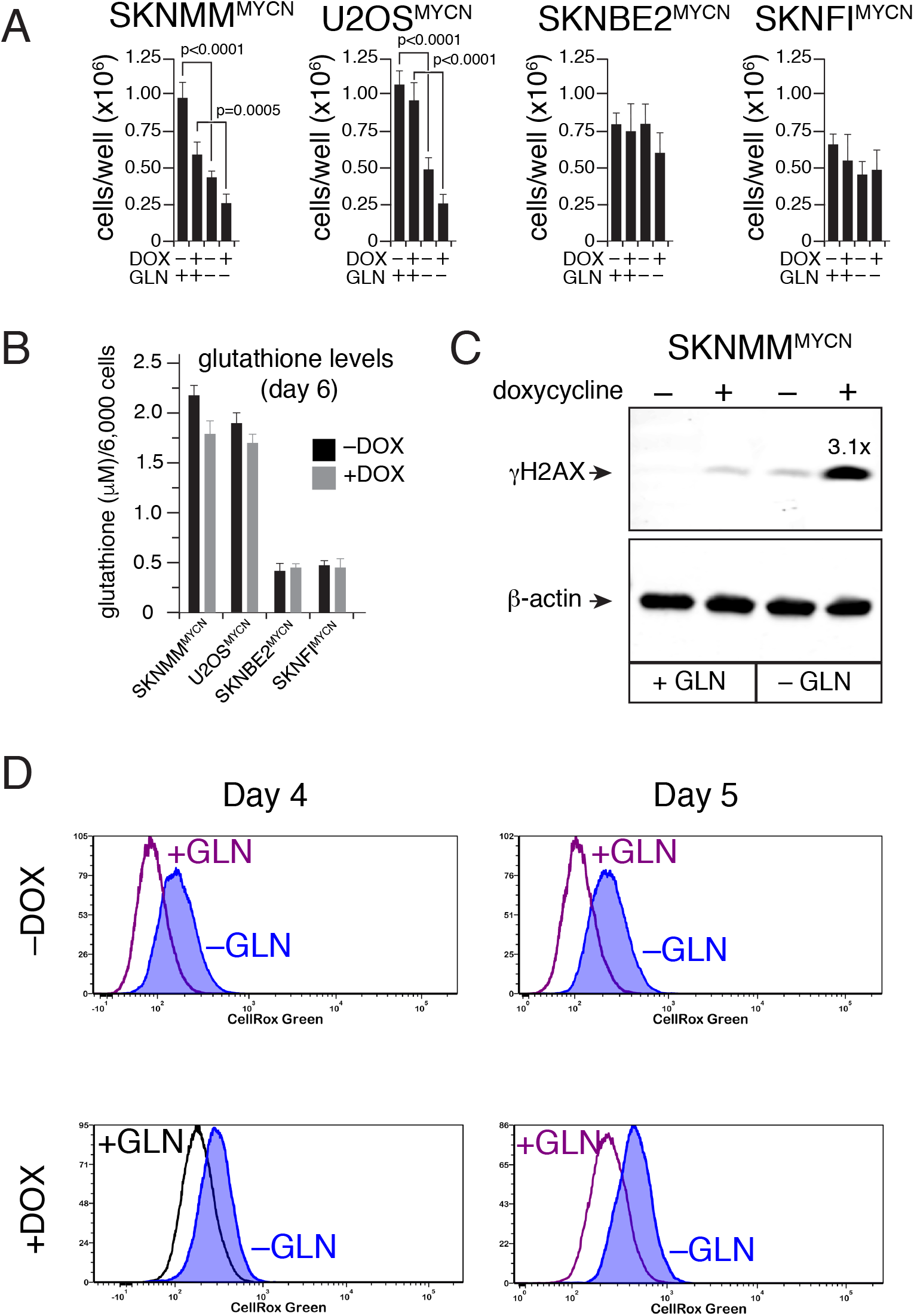
Glutamine metabolism following MYCN induction. **A**) Bar plot of the scoring of the number of cells per well for each cell line cultured in the presence or absence of glutamine with and without doxycycline. Each bar is the mean and standard deviation of triplicate wells. **B**) Bar plot of glutathione levels in the cells on day 6 in culture for each cell line in the presence and absence of doxycycline. **C**) Immunoblot of γ-H2AX and β-actin in the presence and absence of doxycycline and glutamine (GLN). The fold induction is indicated over the γ-H2AX band in the presence of doxycycline relative to the absence of doxycycline in the absence of glutamine. **D**) Flow cytometry of cells stained with CellRox Green to measure reactive oxygen species (ROS) on day 4 and day 5 in the presence and absence of doxycycline and glutamine.

**Figure S7.**
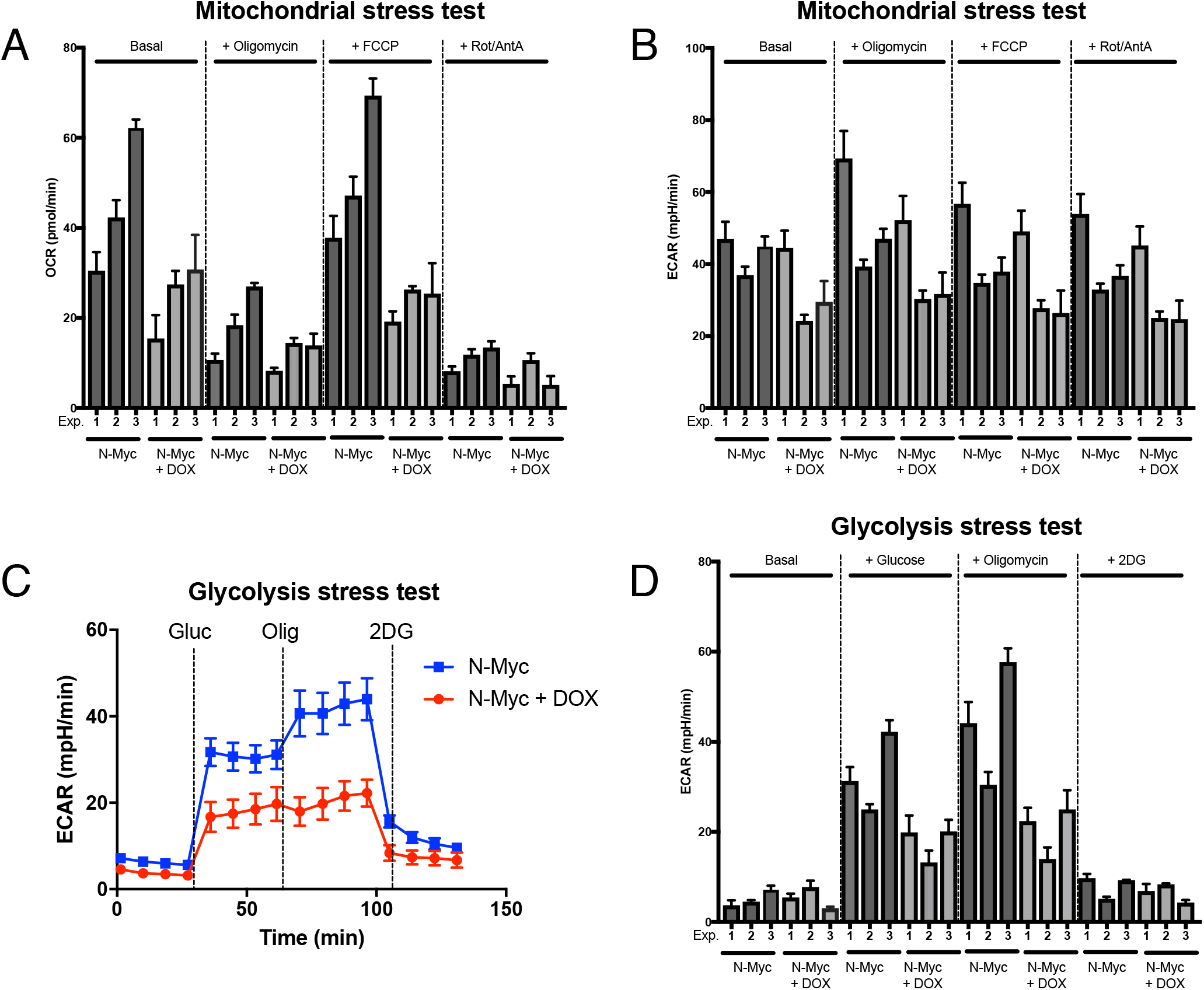
Mitochondrial stress and glycolysis stress for SKNMM^MYCN^ cells. **A**) Bar plot of 3 independent experiments showing the mean and standard deviation for each experiment and each sample at basal and treated conditions. The oxygen consumption rate is plotted in this panel. **B**) The extracellular acidification rate (ECAR) is plotted for the same samples as shown in (A). For the data in A and B, the experiment was performed using conditions for mitochondrial stress testing on the Seahorse instrument. **C**) Representative experiment showing the ECAR for SKNMM^MYCN^ cells in the presence and absence of doxycycline grown under glycolysis stress conditions. **D**) Bar plot of 3 independent experiments for the glycolysis stress test conditions for the indicated cell lines.

**Figure S8.**
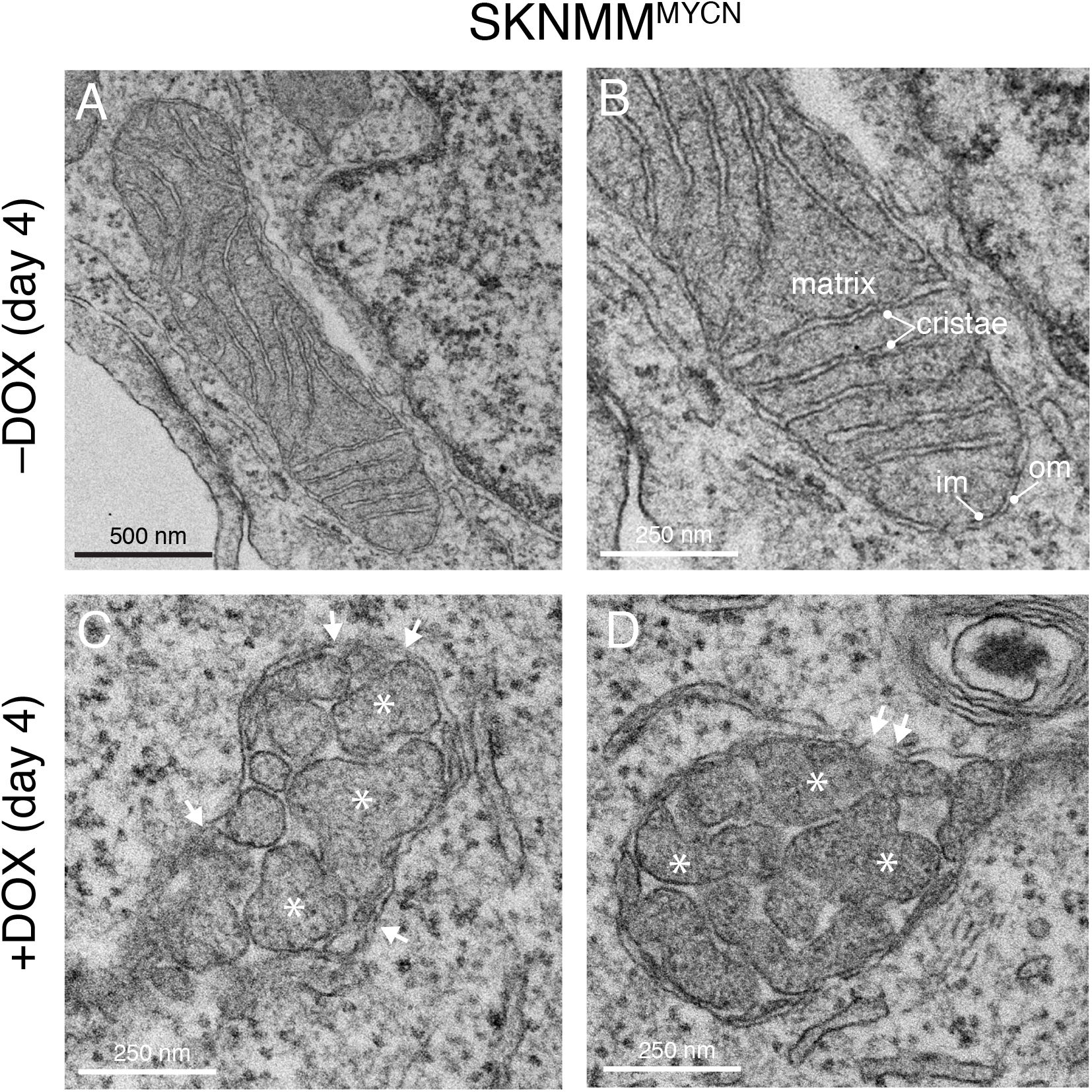
Electron microscopy of mitochondria in SKNMM^MYCN^ cells. **A,B**) Micrograph of SKNMM^MYCN^ cell mitochondria after 4 days in culture in the absence of doxycycline. The mitochondrial matrix and cristae are labeled as well as the inner and outer mitochondrial membranes (im and om). **C,D**) Micrograph of SKNMM^MYCN^ cell mitochondria after 4 days in culture in the presence of doxycycline. The cristae are disorganized and swollen and the mitochondrial membranes are disrupted (arrows).

**Figure S9.**
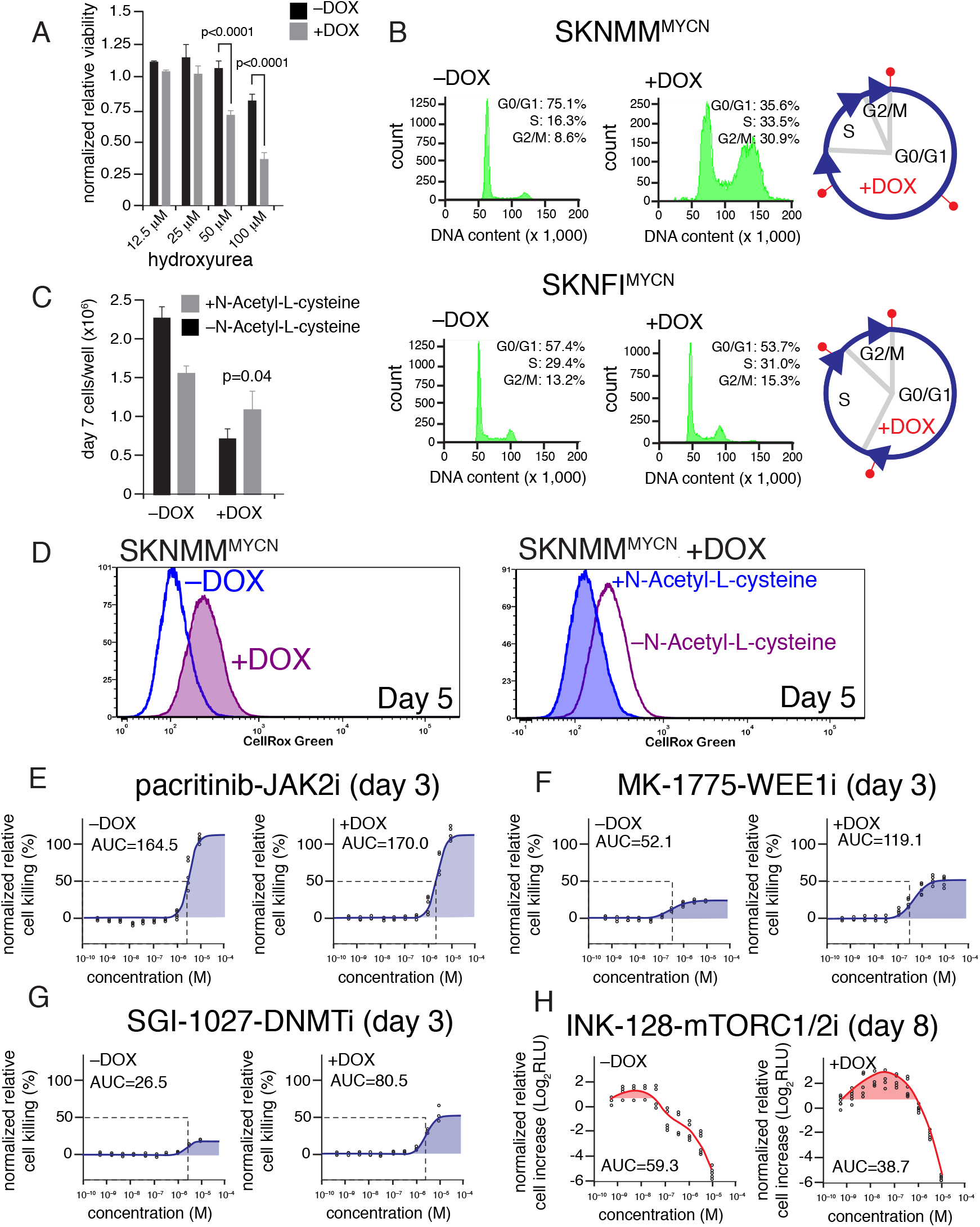
Oxidative stress contributes to synthetic lethality. **A**) Bar plot of the relative viability of SKNMM^MYCN^ cells in the presence and absence of doxycycline on day 4 in culture in the presence of different concentrations of hydroxyurea. Each bar is the mean and standard deviation of triplicate samples. **B**) Plot of DNA content per cell and relative proportion of cells in G0/G1, S and G2/M phases of the cell cycle on day 4 in culture for SKNMM^MYCN^ and SKNFI^MYCN^ cells in the presence and absence of doxycycline. The piechart shows the proportion of each phase for the untreated (-DOX, gray lines) and in the presence of doxycycline (red). **C**) Bar plot of mean and standard deviation of triplicate samples of SKNMM^MYCN^ cells on day 7 after addition of doxycycline in the presence and absence of N-Acetyl-L-cysteine. **D**) The same samples analyzed in (C) were also analyzed by flow cytometry using CellRox Green to measure reactive oxygen species (ROS). The panel on the left shows +/− doxycycline and the panel on the right shows +/− N-Acetyl-L-cysteine in the presence of doxycycline. **E-G**) Representative dose response curves for SKNMM cells on day 3 of culture in the presence and absence of doxycycline for a drug that had no effect on viability (F) and two drugs that potentiated killing my MYCN. The dashed line indicates the EC50 and the area under the curve is shown for each graph. **H**) Spline plot (see Supplemental Information) for one of two drugs that partially rescued the cell death induced by MYCN in SKNMM cells.

## SUPPLEMENTAL MATERIALS AND METHODS

### Patient Samples

Patients were eligible for inclusion in the analytic cohort if they enrolled in the COG Neuroblastoma Biology study ANBL00B1 before treatment; had a confirmed diagnosis of neuroblastoma; and had reported outcome data. Informed consent of the patients and/or parents/legal guardians was obtained at the time of enrollment to ANBL00B1.

### Statistical Analysis of Association of *ATRX* Mutations with Clinical Features

The cohort used in this analysis consists of 477 neuroblastoma patients (of which 476 had clinical data and 475 had outcome data available) representing a mixture of risk levels, disease stages, and ages at diagnosis. All are enrolled on COG protocol ANBL12B8 (“Analysis of *ATRX* Mutations in Neuroblastoma”) and were assayed for *ATRX* mutations. Roughly 80% of the patients were diagnosed between 2010 and 2012, with the remaining 20% diagnosed between 2001 and 2009.

The frequency of *ATRX* mutations, along with Clopper-Pearson exact 95% confidence intervals, were calculated across the entire cohort and in subgroups formed by the combination of age categories (12yrs at diagnosis) and INSS stage categories (1, 2A/B or 3, 4S, and 4). Chi-square tests or Fisher’s exact tests, depending on sample size, were used to test for associations between *ATRX* mutations and clinical factors. Event-free survival (EFS) and overall survival (OS) were compared between patients with vs. without *ATRX* mutations, across the entire cohort and in subgroups based on stage, age, and risk. EFS and OS Cox models were used to assess the prognostic strength of *ATRX* mutations in the presence of standard clinical risk factors.

A forward-selection process was used to construct parsimonious Cox models. At the beginning of the forward-selection process, the following candidate variables were considered to be on equal footing: age at diagnosis (<18months vs. ≥18 months) INSS stage (non-stage 4 vs. stage 4), sex, *MYCN* status (non-amplified vs. amplified), ALK mutation, ploidy (hyperdiploid vs. diploid), 1p (no loss of heterozygosity (LOH) vs. LOH), 11q (no LOH vs. LOH), INPC histology (favorable vs. unfavorable), *ATRX* mutation, MKI (low/intermediate vs. high), and grade (totally undifferentiated/poorly differentiated vs. differentiating). Variables were entered into the model one at a time, with the variable chosen for entry being the one that is most significant at each step, per a Wald test. If at any point in the process histology was chosen to enter the model, then MKI, grade, and age at diagnosis were no longer considered for entry, since histology is confounded with these variables. Conversely, if at any point MKI, grade, or age at diagnosis entered the model, then histology was no longer considered for entry. The selection process ended when all remaining candidate variables failed to reach significance at the 0.05 level.

EFS time was measured as the number of days from diagnosis to date of relapse, disease progression, secondary malignancy, death, or, if no event occurred, date of last contact. OS time was measured as the number of days from diagnosis to date of death, or, if the patient did not die, date of last contact. *MYCN* amplification status was assessed both by the neuroblastoma reference lab as part of ANBL00B1 enrollment and by St. Jude Children’s Research Hospital. There were some discrepancies between the two, particularly in one patient with an *ATRX* mutation (an aberration which previous research suggests is mutually exclusive from MYCN amplification), whom the 6 neuroblastoma reference lab determined to have MYCN amplification, whereas St. Jude detected no amplification. Due to these discrepancies, both sets of MYCN amplification data will be considered individually in this analysis.

Outcome was not compared in the <18 months old at diagnosis subgroup since there were no patients with *ATRX* mutations in that subgroup. Nor was outcome compared in the 18mo-5yrs old at diagnosis subgroup since only one patient in that subgroup had an *ATRX* mutation. The frequency of *ATRX* mutations among the entire cohort of 477 was 4%, with a 95% confidence interval of (2.4%, 6.2%). The highest mutation frequency was observed in the subgroup of INSS stage 4 patients, aged 5-12 years at diagnosis, where 24.1% (13.5%, 37.6%) had some kind of *ATRX* mutation. The lowest mutation frequency was observed in the subgroup of INSS stage 1, 2A/B, or 3 patients, aged 18 months to 5 years at diagnosis, where only 1.3% (0.03%, 7.1%) had a mutation. These results are consistent with the previous finding of *ATRX* mutations being more common among older patients.

Significantly higher mutation frequencies were detected among patients with the following characteristics: INSS stage 4 (vs. non-stage 4), high risk (vs. low/intermediate risk), 11q LOH (vs. no LOH), unfavorable histology (vs. favorable histology), and ≥18 months old at diagnosis (vs. <18 months old). Significant differences in the frequency of mutations were also detected between racial categories (White, Black, Asian, and other) and age groups (12yrs).

The median follow-up time for patients who did not have an event was 3.9 years. The median follow-up time for patients who did not die was also 3.9 years. The estimated 4-year EFS rate and 95% CI for the entire cohort was 81.4% (76.2%, 86.5%) and the estimated 4-year OS rate and 95% CI was 88.6% (84.5%, 92.8%). Across the entire cohort, patients without *ATRX* mutations fared significantly better in terms of both EFS and OS compared to those with mutations (EFS: p12 years old at diagnosis (p=0.0038). Only in the non-INSS stage 4 subgroup did the presence of *ATRX* mutations significantly affect OS, where those without mutations fared significantly better (p=0.0042).

The presence of *ATRX* mutations had a significant effect on outcome in both EFS and OS Cox models. Compared to those without a mutation, patients with an *ATRX* mutation were estimated to have 5.053 times the hazard of an EFS event (p<0.0001) and 3.432 times the hazard of death (p=0.0046). The forward-selection process substituting MYCN amplification data from Dr. Federico’s lab for those from the NBL reference lab yielded the same final EFS and OS Cox models. The effect of *ATRX* mutations on outcome was also tested among patients with INSS stage 4 disease who were at or above 5 years of age at the time of diagnosis. This subgroup was of interest because of its association with a chronic/indolent course of disease. No significant differences in either EFS or OS were detected between patients with vs. without *ATRX* mutations in this subgroup. A notable difference between this subgroup and the subgroups in which significant associations between outcome and *ATRX* mutation were observed, was the number of patients with high-risk disease; the subgroup of older INSS stage 4 patients was entirely highrisk, whereas the other subgroups included a mixture of risk levels. This prompted us to perform a subgroup analysis based on risk group (high vs. low/intermediate). 7 Per log-rank testing, the presence of *ATRX* mutations was not significantly associated with EFS nor OS among patients with high-risk disease, but was significantly associated with outcome for patients with low/intermediate risk disease. There were 5 low/intermediate risk patients with *ATRX* mutations, two of which experienced an event and one subsequently died. Although there was a small number of patients and only a few events, the significant log-rank test results in this subgroup (p=0.0021 for EFS and p=0.0226 for OS) indicate that it is unlikely that those events would have occurred as early as they did, or at all, if there was no association between *ATRX* mutations and outcome among low/intermediate risk patients. The differential effect of *ATRX* mutations between high- and non-high-risk disease groups was also detected in the Cox model of EFS that included an interaction term for mutation and risk level. This model indicated that *ATRX* mutations significantly increased the hazard of an EFS for low/intermediate risk patients, but may have had no effect for high-risk patients. The corresponding OS Cox model did not indicate a significant effect of *ATRX* mutations nor a significant interaction with risk level.

To test the mutual exclusivity between ATRX-mutations and *MYCN* amplification in neuroblastoma, we merged the COG, TARGET, and PCGP data.

Below are results of the analysis looking at the relationship between *ATRX* and *MYCN* in the subgroup of patients with Stage 4, >5 years of age:

| *Variable* | *N (%)* | *ATRX mutation (%)* | *p-value* |
| --- | --- | --- | --- |
| ***MYCN (investigator data)*** |  |  |  |
| *Not amplified* | 153 (90.0) | 41 (26.8) | 0.0135 |
| *Amplified* | 17 (10.0) | 0 (0) |  |

If we look just at Stage 4, 5-12 years of age:

| *Variable* | *N (%)* | *ATRX mutation (%)* | *p-value* |
| --- | --- | --- | --- |
| ***MYCN (investigator data)*** |  |  |  |
| *Not amplified* | 113 (88.3) | 25 (22.1) | 0.0414 |
| *Amplified* | 15 (11.7) | 0 (0) |  |

We also looked at stratified analyses using the Cochrane-Mantel-Haenszel test. Stratifying/controlling by age (<18 months, 18 months-5 years, 5-12 years, >12 years), a significant difference was found in the proportion of *ATRX* mutations in the MYCN nonamplified vs. amplified groups, using *MYCN* combined COG, TARGET, PCGP data (p=0.0037).

### *ATRX* mutations by *MYCN* – Stratified by Age

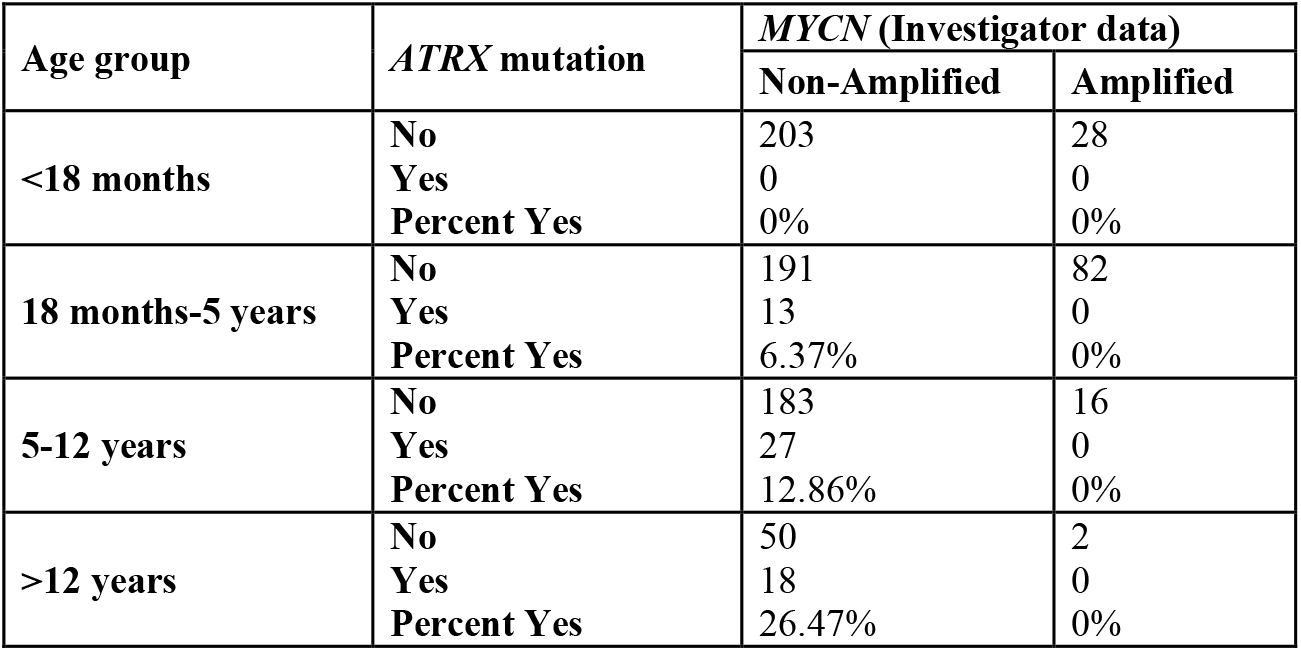

Stratifying/controlling by stage (non-Stage 4, Stage 4), a significant difference was found in the proportion of *ATRX* mutations in the *MYCN* non-amplified vs. amplified groups, using *MYCN* investigator data (p<0.0001).

### *ATRX* mutations by *MYCN* – Stratified by Stage

| Stage group | *ATRX* mutation | *MYCN* (Investigator data) |  |
| --- | --- | --- | --- |
|  |  | Non-Amplified | Amplified |
| Non-Stage 4 | No | 304 | 3 |
|  | Yes | 5 | 0 |
|  | Percent Yes | 1.62% | 0% |
| Stage 4 | No | 323 | 125 |
|  | Yes | 53 | 0 |
|  | Percent Yes | 14.1% | 0% |

Stratifying/controlling by age*stage (non-Stage 4, 18 months-5 years; Stage 4, 18 months-5 years; non-Stage 4, >5 years; Stage 4, >5 years), a significant difference was found in the proportion of ATRX mutations in the *MYCN* non-amplified vs. amplified groups, using *MYCN* COG, TARGET, PCGP data (p=0.0002).

### *ATRX* mutations by *MYCN* – Stratified by Stage*Age

| Stage*Age group | *ATRX* mutation | *MYCN* (Investigator data) |  |
| --- | --- | --- | --- |
|  |  | Non-Amplified | Amplified |
| Non-Stage 4,<br>18 months-5 years | No | 73 | 2 |
|  | Yes | 1 | 0 |
|  | Percent Yes | 1.35% | 0% |
| Stage 4,<br>18 months-5 years | No | 118 | 80 |
|  | Yes | 12 | 0 |
|  | Percent Yes | 9.23% | 0% |
| Non-Stage 4,<br>>5 years | No | 121 | 1 |
|  | Yes | 4 | 0 |
|  | Percent Yes | 3.2% | 0% |
| Stage 4,<br>>5 years | No | 112 | 17 |
|  | Yes | 41 | 0 |
|  | Percent Yes | 26.8% | 0% |

### Custom Capture for *ATRX, MYCN* and *ARID1A/B*

Targeted enrichment was performed using the Seqcap EZchoice Kit (Roche) according to vendor instructions for the Kapa workflow with 500 ng of genomic DNA as the starting input for library construction.

### Exome Sequencing

Whole exome sequencing was conducted using the SeqCap EZ HGSC VCRome (Roche) according to manufactures instructions.

## MYCN FISH

Purified human *NMYC* BAC DNA (RP11-1183P10) was labeled with a red-dUTP (AF594, Molecular Probes) and human chromosome 2 control (2q11.2) BAC DNA (RP11-527J8) was labeled with a green-dUTP (AF488, Molecular Probes) both by nick translation. The paraffin slides were deparaffinized with xylene 2x 10 minutes each at RT, placed in ETOH 3x 2 min each at RT, air-dried, placed in 10% buffered formalin for 1 hour at RT. The slides were then placed in pepsin (8mg/ml) in 0.1N HCL for 3 minutes, rinsed in dH20 for 5 min at RT. One-hour incubation in Carnoy’s fixative at RT was performed. The labeled probes were combined with human sheared DNA and hybridized to the slides in a solution containing 50% formamide, 10% dextran sulfate, and 2X SSC. The probe and slide were co-denatured at 90°C for 12 min and incubated overnight at 37°C on a Thermobrite. Briefly washed in PN and then stained with DAPI (1μg/ml). Images were captured using a Plan-Apochromat 63X objective on a Zeiss Axio Imager.Z2 microscope and GenASIs scanner (ScanView software version 7.2.7).

## ATRX IHC

ATRX immunohistochemistry was done on formalin-fixed 4-μm-thick paraffin sections using anti-ATRX antibodies polyclonal antibodies (1:600; Sigma-Aldrich) as previously described ^1^.

### Orthotopic Patient Derived Xenografts

Mouse studies were performed in a strict accordance with the recommendations in the Guide to Care and Use of Laboratory Animals of the National Institutes of Health. The protocol was approved by the Institutional Animal Care and Use Committee (IACUC) at St. Jude Children’s Research Hospital. All efforts were made to minimize suffering. All mice were housed in accordance with approved IACUC protocols. Animals were housed on a 12–12 light cycle (light on 6:00, off 18:00) and provided food and water ad libitum.

Patient-derived xenograft cells were maintained orthotopically in athymic nude (Jackson Laboratories, strain code 007850), NSG (Jackson Laboratories, strain code 005557) or C57BL/6 scid mice (Jackson Laboratories, strain code 001913) as described before (Stewart et al, 2017). Tumor cells were dissociated using Trypsin (10 mg/ ml, Sigma Cat # T9935) filtered and then suspended in a Matrigel basement membrane matrix (BD Biosicences catalogue number 354234) at a concentration of 2×10^5^ cells per 10-μl and placed on ice for injection. The cells were injected in the para-adrenal region of the mice with ultrasound guidance under anesthesia.

### Genetic mouse models

To test the incompatibility between *ATRX* mutations and *MYCN* overexpression in mice, we used *LSL-MYCN:DbhiCre* mice^2^. These mice express *MYCN* from the *Rosa26* locus when a stop sequence is floxed out by Cre recombinase under the control of *Dbh* promoter in sympathetic ganglion cells. We breed *LSL-MYCN:DbhiCre* mice to *Atrx^Flox^* mice^3^ to inactivate *Atrx* in the same tissues. We also tested the consequences of *Atrx* inactivation when *Th-ALKF^1174L^* allele is added^4^ (*LSL-MYCN: DbhiCre:Th-ALKF^1174L^*). *Th-ALK^F1174L^* mice constitutively express *ALK^F1174L^*, the most common activating *ALK* mutation found in human neuroblastoma, under the control of *Th* promoter.

Mice were genotyped using the published protocols^2–5^. Mice with the desired genotypes were enrolled in the study at the time of weaning and were followed up by abdominal palpation at least once every 2 weeks starting from age of 6 weeks. Enrolled mice that died with no detectable tumors were censored out of the survival study. Mice were euthanized when the tumor is 20% of body weight or when the mouse has other morbidities e.g. lethargic, hunched or dehydrated. After euthanasia, mice were opened longitudinally to expose abdomen, thorax and neck to search for tumors. Tumors were dissected out and were cut into pieces for histological, immunohistochemical, DNA, RNA and protein preparations.

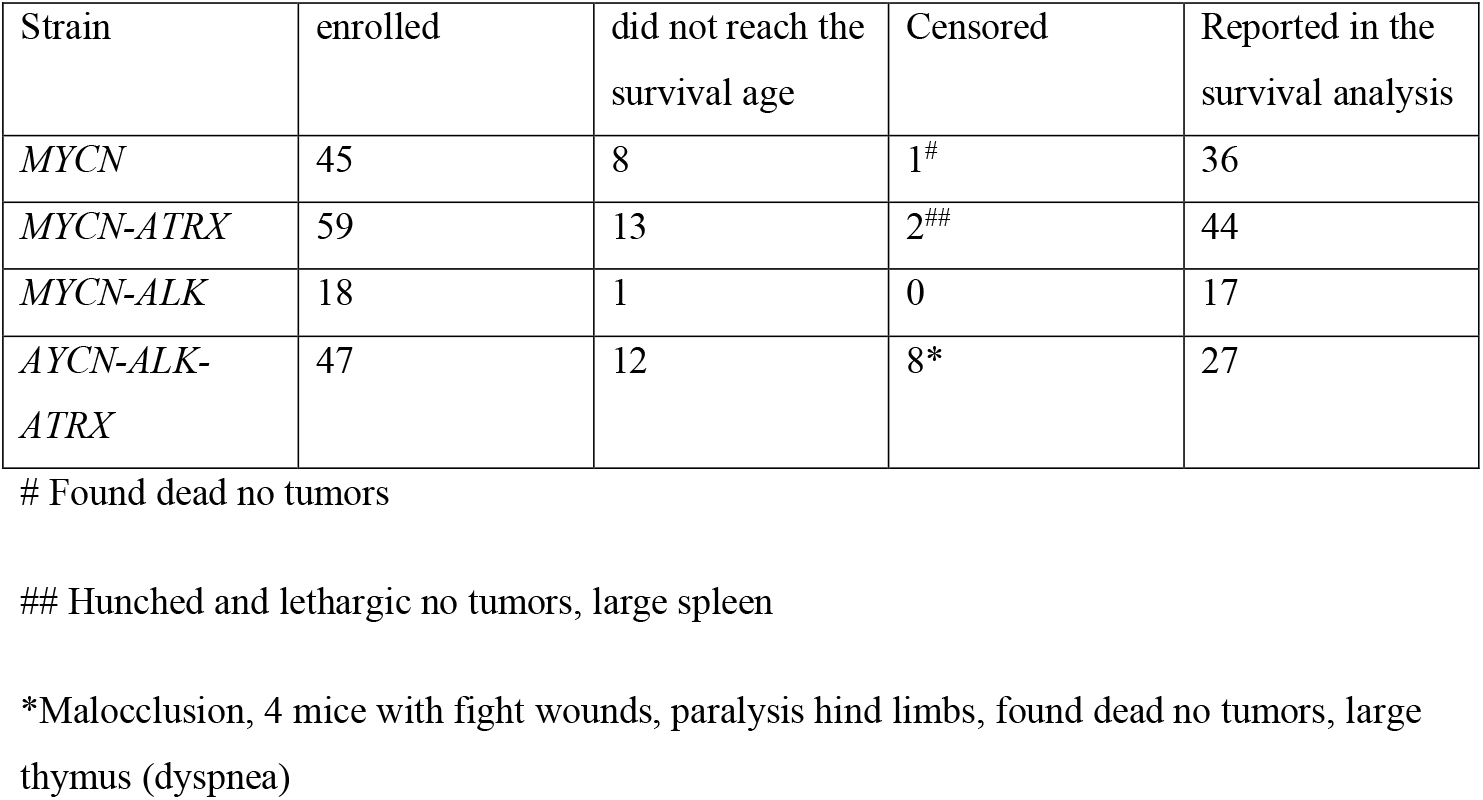

| Strain | enrolled | did not reach the survival age | Censored | Reported in the survival analysis |
| --- | --- | --- | --- | --- |
| *MYCN* | 45 | 8 | 1<sup>#</sup> | 36 |
| *MYCN-ATRX* | 59 | 13 | 2<sup>##</sup> | 44 |
| *MYCN-ALK* | 18 | 1 | 0 | 17 |
| *MYCN-ALK-ATRX* | 47 | 12 | 8* | 27 |

### Whole Genome Sequencing

DNA from the tumors and matching germline samples was sequenced and analyzed as described previously^6^. Mutations, sequence variants, copy number alterations and structural variations were detected as previously described^7^. WGS data were uploaded at EMBL with accession number EGAS00001002528.

### Whole Genome Bisulfite Sequencing and analysis

Genomic DNA was extracted using the DNeasy Kit (QIAGEN #69504) according to the manufacturer’s protocol and quantified with a Nanodrop (Thermoscientific). Whole genomic bisulfite conversion and sequencing were done by Hudson Alpha (Huntsville, Al).

Sequencing data were aligned to the hg19 human genome using BSMAP2.74^8^. We first performed statistical test of differentially methylated locus (DML) between MYCN-amplified and non-MYCN-amplified samples using DMLtest function (smoothing=TRUE) in DSS (Wu et al 2015), the results were then used to detect differentially methylated regions (DMRs) using CallDMR function in DSS, the p-value threshold for calling DMR is 0.01. The minimum length for DMR is 50 bps and the minimum number of CpG sites for DMR is 3. The minimum methylation difference is 0.2. The DMRs that overlap with the promoters (1 kb flanking the TSS) of differentially expressed genes (at least 2-fold changes and adjust p value <=0.1) were deemed to be associated with gene expressions.

### RNA-Seq and analysis

RNA was extracted from cultured cells and tissues using RNeasy Kit (QIAGEN #74104) or TRIzol (Life Technologies) preparations via a phenol–chloroform extraction or Direct-zol™ RNA MiniPrep (Zymo Research # R2052) following manufacturers’ protocols. RNA concentration was measured using a NanoDrop (Thermo Fisher scientific, Waltham, MA) and the quality of RNA was determined with a bioanalyzer (Agilent Technologies, Santa Clara, CA). Libraries were prepared using the TruSeq Stranded Total RNA Library Prep Kit (Illumina, San Diego, CA) from 500-ng total RNA. Paired end deep sequencing was done using HiSeq 2500 sequencers (Illumina, San Diego, CA).

FASTQ sequences were mapped to the human genome hg19 (GRCh37-lite) and Gene-based FPKM quantification was calculated similar to previously described^9^ using GENCODE gene annotation (v26lift37). We then used VOOM^10^ for differential analysis between +DOX vs -DOX. To confirm the results were meaningful, we ranked the genes by log2 fold change and use GSEA^11^ to check if any MYCN related gene sets in MSigDB(v6.1) could be found enriched. Indeed, WEI_MYCN_TARGETS_WITH_E_BOX^12^ gene set were found top enriched for up-regulated genes in +DOX (rank 46 out of 11371 gene sets, NES=3.01, FDR corrected p-value < 1e-4).

### ChIP-Seq

Antibodies used in this study to generate ChromMM were previously validated and the validation data and protocols are available through St. Jude Childhood Solid Tumor Network website (https://www.stjude.org/CSTN/)^13^.

For MYCN ChIP-Seq, we used rabbit polyclonal antibody (Active Motif Cat# 61185). This antibody was validated by small-scale and large-scale sequencing by Active Motif as well as by us in St. Jude Children’s Research Hospital. We tried different amounts of the antibodies and chromatin and determined that 20-ug of the antibody and 200-ug of the chromatin per ChIP-Seq reaction show the best results.

Chromatin in cultured cells was fixed by changing the medium with fresh medium containing 1% formaldehyde (Thermo scientific # 28906). Snap-freezed xenografts were pulverized and hammered to powder in liquid Nitrogen, suspended in PBS and fixed by adding formaldehyde (Thermo scientific # 28906) to final concentration of 1%. Fixation was done for 10 minutes at room temperature. Cross linking was stopped by adding glycine to a final concentration of 1.25 M. Cells and xenografts were washed in 1X PBS containing 1X protease inhibitor (Pierce # 78430).

Chromatin preparation and shearing were done using TruChIP chromatin shearing kit (Covaris #520127) following the manufacturer’s protocol. Chromatin immune-precipitation (ChIP) reactions were performed using iDeal ChIP-seq kit (Diagenode # C01010051) with 10% of chromatin from each reaction was used as an input. Precipitated DNA was de-crosslinked, extracted using MinElute PCR-purification kit (QIAGEN #28006) and quantified using the Quant-iT PicoGreen ds DNA assay (Life Technologies #Q33120). Quantitative polymerase chain reaction (qPCR) was done to assess the efficiency of ChIP reactions as described in the validation protocols. Five to ten nanograms of the input and precipitated DNA were used to prepare the sequencing libraries using the NEBNext ChIP-Seq Library Prep Reagent Set for Illumina with NEBNext High-Fidelity 2 × PCR Master Mix following the manufacturer’s instructions (New England Biolabs) with some modifications as described previously (Aldiri et al 2017); after adaptor ligation, 1:1 Ampure cleanup was added twice, no Ampure size-selection step and the extension was done for 45 seconds. Inset-size distribution of the completed libraries was analyzed using a 2100 BioAnalyzer High Sensitivity kit (Agilent Technologies) or Caliper LabChip GX DNA High Sensitivity Reagent Kit (PerkinElmer). ChIP reactions and library preparation for RNA-Pol II, Brd4 H3K9me3 and MYCN were done by Active Motif. Quant-iT PicoGreen ds DNA assay (Life Technologies) and Kapa Library Quantificaiton kit (Kapa Biosystems) or low-pass sequencing on a MiSeq nano kit (Illumina) were used to quantify the libraries. Fifty-cycle single-end sequencing was performed on an Illumlina HiSeq 2500.

### ChIP Seq analysis

ChIP-Seq analysis was done as described previously^14^. Briefly, ChIP-Seq reads were aligned to human genome hg19 (GRCh37-lite) using BWA software (version 0.7.12-r1039, default parameter) and the duplicated reads were marked using Picard software. We kept only nonduplicated reads for the analysis using samtools (parameter “-q 1 -F 1024” version 1.2). The quality control of the data followed ENCODE criteria. We calculated relative strand correlation value (RSC) and estimated the fragment size under support of R (version 2.14.0) with packages caTools (version 1.17) and bitops (version 1.0-6). We required at least 10M unique reads for point source factors H3K4me2/3, H3K9/14Ac, H3K27Ac, CTCF, RNAPolII, BRD4) with RSC > 1, 20M unique reads for broad markers (H3K4me1, H3K9me3, H3K27me3, H3K36me3) and 10M unique mapped reads for INPUTs with RSC < 1. All samples were manually inspected and the SPP (version 1.1) was used to generate the cross-correlation plot. Then we generated bigwig files from the best fragment size (the smallest fragment size estimated by SPP). Bigwig files were examined using IGV genome browser for clear peaks and low background noise. MACS2(version 2.0.10.20131216)^15^ were used to call peaks for MYCN ChIP-Seq data with FDR corrected p-value 0.05(--nomodel –extsize FRAGMENTSIZE, where FRAGMENTSIZE was estimated by SPP as described above). Genes promoter (TSS +/− 2kb) with MYCN peaks called were considering have MYCN binding site. Enrichr^16^ were used for pathway analysis.

### Super-Enhancer and core regulator circuit analysis

Super-Enhancers and core regulator circuit analysis have been performed as described previously^14^. We called auto-regulators by CRCMapper^17^ based on Super-Enhancers called by ROSE^18^. We provided two sets of results that one only used H3K27ac data while the other one excluded H3K27ac peaks overlapping promoter (defined by H3K4me3 peaks) before calling Super-Enhancer by ROSE.

### ChromHMM

ChromHMM models were generated for y as described before^6^. We applied the Hidden Markov Model trained for 18 states from previously 4 types of solid tumors to the data from this manuscript. Expanded chromHMM state 13 were defined as proportion of promoter downstream (TSS ~ TSS+2kb) in SKNMM^MYCN^ +Dox > 2 fold of SKNMM^MYCN^ −DOX and >= 20% in SKNMM^MYCN^+DOX.

### Immunoblotting

The medium was removed and the cells were washed twice with phosphate-buffered saline (PBS) then the cells were scrapped in 1-ml PBS and centrifuged at 2000Xg for 5 minutes at 4°C to pellet the cells. PBS was removed and the cells were flash-freezed in dry ice and stored at - 80°C until use. Cell pellets were thawed on ice and lysed for 15 min in RIPA buffer (Sigma Cat. #R0278) supplemented with 1 X Halt protease inhibitor (ThermoFischer, Cat. #7834) and 5 mM EDTA. Lysate were then centrifuged for 15 minutes at 15,000 rpm at 4°C. Protein concentration in the cell lysates was measured using BCA protein assay kit (ThermoFischer, Cat. # 232225). Equal amounts of total protein were resolved on 4-15% SDS-PAGE gel (Biorad Cat#4561086) and transferred to a nitrocellulose membrane by electrophoresis at 30 volts overnight at 4°C. Non-specific binding was blocked by incubating the membrane with Odyssey Blocking buffer, PBS (LiCore Cat# 927-40000) for 1 hour at room temperature. Then the membrane was incubated with the primary antibody overnight at 4°C. The following antibodies were used: rabbit anti-ATRX (1:1000, Abcam, ab97508), mouse anti-MYCN (1:1000, Santa cruz Biotechnology, sc-53993), rabbit anti-GAPDH (1: 2,500, Abcam ab-9485), rabbit anti-DAXX (1:1000, Santa cruz Biotechnology, sc-7152) and mouse anti- -actin (1: 5,000, Sigma, A1978). Membranes were washed three times in PBS with 0.1% Tween-20 at room temperature then were incubated in the secondary antibodies: IRDye 800CW goat anti-mouse (1: 5,000, LiCore, 925-68070) or IRDye 680CW goat anti-rabbit (1: 5,000, LiCore, 925-32211) for 30 minutes at room temperature. The membranes were then washed three times in PBS-T; 5 minutes each at room temperatures and one time in PBS. The membranes were then scanned by Odyssey CLX infrared gel imaging system (Li-Cor Biosciences, Lincoln, NE). Band intensities were quantified using ImageStudio Lite software (Li-Cor Biosciences, Lincoln, NE).

### Telomere FISH

After a 5 hour colcemid incubation, cells from both samples were harvested using routine cytogenetic methods. A commercially prepared directly FITC labeled PNA telomere probe (DAKO, Cat. # 5327) was used for this analysis. DAKO protocols were followed for the pretreatment and hybridization steps. The slides were then stained with 4,6-diamidino-2-phenylindole (DAPI). Two hundred fifty-five interphase nuclei from each sample were individually analyzed for the number of telomeres/cell, fluorescence/cell, and fluorescence/telomere. A fixed exposure time is used for all cells.

### Telomere qPCR

Telomere quantitative polymerase chain reaction (QPCR) was done as described previously (Cheung et al. 2012). DNA was extracted using DNAeasy Kit (Qiagen, Cat. # 69504). qPCR was done on 10-ng DNA template using SYBR green Select master mix (Thermo Fisher, Cat. # 4472908). Two sets of primer pairs were used in separate reactions were used to amplify the telomeric sequence and a common locus RPLP0 which was used as an internal control. The reactions were done in duplicates and the average delta C(t) was calculated for every sample by subtracting the average C(t) values of the telomeric reaction from those of the internal control reactions. The primers used are

> Telomere F: 5-CGGTTTGTTTGGGTTTGGGTTTGGGTTTGGGTTTGGGTT-3
>
> Telomere R: 5-GGCTTGCCTTACCCTTACCCTTACCCTTACCCTTACCC-3

Control, locus:

> RPL0 F: 5-CAGCAAGTGGGAAGGTGTAATCC-3
>
> RPL0 R: 5-CCCATTCTATCATCAACGGGTACAA-3

### *ATRX* shRNAs

Eight ATRX-specific shRNA lentiviral vectors were purchased from Dharmacon as bacterial glycerol stocks (Cat. # V2LHS-092920 (#9), V2LHS-202920 (#20), V2LHS-22887 (#22), V3LHS-375686 (#37), V2THS-22887 (#87), V2THS-93289 (#30), V2THS-255090 (#13) and V2THS-201895 (#95)). Lentiviruses were made in HEK293T cells by co-transfecting the viral vectors with 3 packaging plasmids. The supernatants containing the viral particles after 48 and 72 hours were collected, filtered, concentrated using ultracentrifugation and titrated in HeLa cells using the GFP or RFP reporters in the vector. In addition, 2 non-silencing shRNA lentiviral particles were used as controls (Cat. # RHS4743 and RHS4346). Cells were transduced with equal number of viral particles and transduced cells were selected with flow cytometry or pouromycin after 48 hours post transduction.

### Cell Cycle Analysis

Cells were trypsinized and harvested, pelleted and suspended in 1-ml propidium iodide solution (0.05 mg/ml propidium iodide (Sigma, Cat. # P4864), 0.1% sodium citrate, 0.1% Triton X-100). The cells were then treated with 10-ml 0.2 mg/ml of ribonuclease A (Colbiochem, Cat. # 556746) for 30 minutes at room temperature, filtered through 40-μm nylon mesh and analyzed by flow cytometry for the DNA content.

### Colony Assays

Cells were transduced with a lentivirus expressing *ATRX* shRNA #9, *ATRX* shRNA #20 or scrambled control shRNA at MOI 5 for 24 hours. Then the cells were harvested, suspended into single cells in a pre-warmed medium, counted and seeded at 300 cells/ well in 6-well plates. The plates were shaken gently every 1-2 minutes for 10 minutes. Fresh medium was added every 4 days. After 2-3 weeks when colonies were visible, cells were washed, fixed in 1% paraformaldehyde for 30 minutes at room temperature and stained with 0.1% Crystal Violet.

### CRISPR/Cas9 Targeting *ATRX* and *DAXX*

Guide RNAs (gRNAs) were designed targeting an early and conserved exon between isoforms of the *hATRX* and *hDAXX* genes. Bioinformatics analysis was performed to identify gRNAs with at least 3bp of mismatch to other sites in the human genome. gRNAs were further validated in human K562 cells for cutting efficiency using targeted deep sequencing.

Two validated *hATRX* (g2 and g5) and one *hDAXX* (g11) gRNAs were used in this study. The sequences of gRNAs are:

> *hATRX.g2*: 5′-CTGCACTGCTTGTGGACAACNGG-3′
>
> *hA TRX.g5*: 5′-CAATGAAGGGTGTCTATAAANGG-3′
>
> *hDAXX.g11*: 5′-GGTTCTTCTGACAGTAACGANGG-3′

Cells were transiently transfected with PC526-Cas-9 plasmid (to express Cas-9; Addgene # 43945), either ZZ52.hATRX.g2 or ZZ52.hATRX.g5 (to express *hATRX* gRNA; the backbone plasmid Addgene # 43860) and pUB-GFP (to express GFP; Addgene # 11155) in a ratio of 7:7:1 respectively using Mirus 2020 (Mirus Biotechnology Cat. # 2020). After 48 hours, the cells were sorted for GFP positive cells. Half of the sorted cells were pelleted immediately and stored at - 80°C and the remaining cells were seeded in 6-well plates and harvested at different time points. To test for the NHEJ efficiency after CRISPR targeting in IMR-32 cells, we transfected the cells with the same mix of plasmids but added a control plasmid expressing gRNA targeting the *hAAVS1* locus (Addgene # 41818) in a ratio of 4:4:6:1 for hAAVS1.gRNA: ATRX.gRNA: Cas-9: GFP expressing-plasmids respectively. DNA was extracted from the cell pellets using DNAeasy kit (Qiagen, Cat. # 69504) and PCR was done to amplify the targeted region using primers to which MiSeq adaptors were annexed and the gel-purified PCR products were subjected to deep sequencing.

The primers used are (adaptor sequences are underlines)

*hATRX* F:

> <u>TCGTCGGCAGCGTCAGATGTGTATAAGAGACAG</u>TGGAATTTGCCAAGGTTGTCATGTGC

*hATRX* R:

> <u>GTCTCGTGGGCTCGGAGATGTGTATAAGAGACAG</u>CTTTCTTCAAGACTGTGCCCTCAAAGGCC

*hAAVS1* F:

> <u>TCGTCGGCAGCGTCAGATGTGTATAAGAGACAG</u>GTGTCCCCGAGCTGGGACCA

*hAAVS1* R:

> <u>GTCTCGTGGGCTCGGAGATGTGTATAAGAGACAG</u>TAGGAAGGAGGAGGCCTAAGGATGGG

The percentage of indels were determined using CRISPResso software as described before^19^.

### Comet assay

Alkaline Comet Assay was done using CometAssay Reagent Kit (Trivegen, Cat. # 4252-40-K) following the manufacturer’s protocol. Briefly, cells were harvested and suspended in PBS buffer at a concentration of 1X10^5^ cells/ml. Cells were combined with LMAgarose in a ratio 1:10 at 37° C and 50-1 of the mix was spread over the CometSlide™. The doxycycline-treated and untreated samples were loaded onto 2 different wells on the same slide. Slides were left at 4°C for 30 minutes in the dark to allow agarose to polymerize. The slides were then incubated in a freshly-made Alkaline Unwinding Solution (200 mM NaOH, 1 mM EDTA, pH> 13) overnight at 4°C in the dark. Electrophoresis was done at 4°C for 30 minutes at 21 volts in the Alkaline Electrophoresis Solution (200 mM NaOH, 1 mM EDTA, pH> 13). The slides were washed twice in dH_2_O, dried at 37 °C and stored in the dark. To analyze the samples, nuclei suspended in the agarose gel were stained by 100- 1 of SYBR^®^ Gold (Thermo Fischer, Cat. # S 11494) diluted 1:30,000 in TE buffer (10 mM Tris-HCL pH 7.5, 1 mM EDTA). Then the slides were rinsed in water and dried completely at 37 °C. Slides were examined by a fluorescence microscopy for excitation/ emission: 496/522 and 10-15 images were taken per sample. Percentage of DNA in the tail was quantified in the images using ImagJ software by measuring fluorescence intensity in the tail/total fluorescence X 100. Mann-Whitney non-parametric statistical test was used to compare the scores in the samples with or without doxycycline.

### Spectral karyotyping (SKY) analysis

Cells were treated with colcemid for 5 hours at 37°C then the cells were harvested for cytogenetic analysis. SKY analysis was done using a commercially available probe from spectral imaging (Carlsbad, CA) following the manufacturer’s protocol. Fifty metaphase cells from every sample were scored for the presence of DNA fragmentations. Statistical analysis was done using Fischer exact test.

### Hydroxyurea sensitivity

The cells were seeded in 96-well plates at a density of 2,500 cells per well over night. Next day (day 0), the medium was changed with a fresh medium with or without 2 g/ml doxycycline. After 24 hours of doxycycline treatment (day 1), the medium was changed again with fresh medium containing serially diluted hydroxyurea with or without doxycycline. At day 4, the cell viability was assessed using the CellTitre-Glo^®^ cell viability assay kit (Promega, Cat. # G7572) using PHERAstar FSX (BMG Labtech, San Diego, California).

### Inducible MYCN

To make the doxycycline-inducible cell lines, we used pLenti4/TO/V5 DEST system (Invitrogene, Cat. # V480-20) following the manufacturer’s protocol. Briefly, cells were transduced with the pLenti4/TO lentivirus to constitutively express the Tet repressor and the cells were selected with 10-μg/ ml blasticidin-HCL (Thermo Fisher, R21001) in the medium.

The cells were then transduced with the V5 DEST-MYCN or V5 DEST-control lentivirus and were selected with 10-μg/ ml zeocin (Thermo Fisher, R25001) in the medium. The V5 DEST-MYCN virus was a gift from the Freeman lab at St. Jude Children’s Research Hospital. The cells were maintained in a selective medium containing 5-μg/ ml blasticidin-HCL and 5-μg/ ml zeocin. We also cloned *MYCN* into another dox-inducible system using Lenti-X Tet-One™ (Takara Cat# 631847) following manufacturer protocol. We confirmed the sequence of the construct by sanger sequencing. Cells were transduced with Lenti-X Tet-One™ lentiviruses containing *MYCN* or with the empty vector. Selection for the cells containing the construct was done using puromycin and the cells were maintained in puromycin-containing media. Induction of *MYCN* expression was done by adding doxycycline to the medium to a final concentration of 1-2 μg/ ml.

### Growth Curve

The cells were plated in 6-well plates at a density of 50,000 cells per well over night. Then the medium over the cells was changed with a fresh medium with or without 1-2-μg/ml doxycycline. Three wells from each of the cell lines were harvested every day and counted for the total number of cells per well using a hemocytometer.

### Live Imaging

Live cell time-lapse imaging experiments for SKNMM^MYCN^ in the presence or absence of doxycycline were performed using a Nikon C2 confocal configured on a TE2000E2 microscope with a 20X 0.8 NA Plan Apo lens (Nikon Instruments Inc., Melville, NY). A 640 nm DPSS lasers was used to simultaneously excite Annexin V-Alexa Fluor 647 and generate the DIC images. During imaging experiments, the cells were maintained at 37°C and 5% CO2, and images were collected every 30 min for 67 hours. Images were acquired and processed using NIS Elements software.

### In vivo Mixing Experiment

SKNMM^MYCN^ and SKNMM^CONT^ cells were labelled with luciferase-YFP using a lentivirus. Two mixed cell pools were prepared in a 1:1 ratio: labelled SKNMM^MYCN^ cells/ unlabeled SKNMM^CONT^ and labelled SKNMM^CONT^ / unlabeled SKNMM^MYCN^. Cell mixes were injected in the para-adrenal region of the mice (4 million cells/ mouse) with ultrasound guidance under anesthesia as explained before. In this experiment, we used NOD.CB17-Prkdc^scid^/J female mice (Jackson laboratories, Stock # 001303). The mice were followed up for the tumor formation using ultrasound scanning every 2 weeks, xenogen signal every week and palpation every week. Mice were enrolled in the study when the tumor was at least 14 mm^3^ in size using the ultrasound scanning. Enrolled mice were randomly assigned to receive either doxycycline (2mg/ml) in the drinking water or continue on regular water.

The labelled/ unlabeled cell mixes were also maintained in culture and the ratio of YFP expressing cells in each cell mix was determined weekly using flow cytometric analysis.

### Lactate Measurements

Lactic acid was measured using Lactate Assay Kit (Sigma Cat. # MAK064) following the manufacturer’s protocol. The absorbance at 570 nm (A570) was measured for a colorimetric detection of the lactate using the microplate reader SpectraMax-M5 (Molecular Devices, Sunnyvale, California).

### GC-MS-mediated analysis of 13C-labelled TCA metabolites

The analytical procedure applied was based on the one previously published^20^. Briefly, cells were cultured in glutamine-free medium (10% FBS, penicillin/streptomycin) supplemented with 4mM of standard glutamine or [U-^13^C]-glutamine (CLM-1822-H); Cambridge Isotope Laboratories, Inc. for 5 hours. Next, cells were trypsinized, washed with ice-cold saline solution, snap-frozen and stored in liquid nitrogen for further processing. Extraction of hydrophilic metabolites was performed using 2 volumes of methanol/water solution (2:1 v/v, HPLC-grade, spiked with 5nM of norvaline as internal standard), homogenization and mixing with 1 volume of chloroform (HPLC-grade). Derivatization with 2% methoxylamine-HCl solution and GC-MS-grade MTBSTFA + 1% t-BDMCS reagent was performed as previously described. Samples were analyzed in split-less mode using Agilent 7890B GC system equipped with a 5977B MSD, Agilent VF5ms, +1m EZ, 60m, 0.25, 0.25 μM column, helium carrier gas and Mass Hunter software package. Chromatographic gradient conditions: separation time 36 min; start at 80oC, 1 min; ramp to 250oC at 7oC/min; ramp to 300oC at 50oC/min and hold for 9 min. MSD settings: source 150oC; quadrupole 150oC; interface 300oC; injector 250oC; EI source 70 eV, EI pressure 1.8 x 10 −5 torr. Signals were acquired using SIM mode conditions as described before (5) as well as using scanning mode (full scan range 50-500 m/z at ~ 2 scans/s (N=3)). Enrichment of C13-labelled isotopomers of analyzed compounds was corrected by natural abundance using “standard glutamine” medium-incubated sample as reference.

### Capillary electrophoresis-time of flight mass spectrometry (CE-TOFMS) for detecting metabolites

MYCN-induced and un-induced SKNMM^MYCN^ cells were grown on either glutamine-free or glucose free completed medium supplemented with [U-^13^C]-glutamine (CLM-1822-H) or [U-^13^C] glucose (CLM-1396); Cambridge Isotope Laboratories, Inc. for 18 hours. Metabolites were extracted from the cells following the protocol provided by Human Metabolome Technology Inc, (Boston, MA). Samples were filtered through through 5-kDa cut-off filter (ULTRAFREE-MC-PLHCC, Human Metabolome Technologies, Yamagata, Japan) and sent to Human Metabolome Technologies Inc. for analysis. We also sent snap-freezed medium from the same samples for metabolomic analysis.

Metabolites were measured in both Cation and Anion modes using Agilent CE-TOFMS system (Agilent Technologies Inc.) and Capillary: Fused silica capillary i.d. 50 μm Å~ 80 cm with the following conditions:

<u>Cationic Metabolites (Cation Mode)</u>:

Run buffer: Cation Buffer Solution (p/n : H3301-1001)

Rinse buffer: Cation Buffer Solution (p/n : H3301-1001)

Sample injection: Pressure injection 50 mbar, 10 sec

CE voltage: Positive, 27 kV

MS ionization: ESI Positive

MS capillary voltage: 4,000 V

MS scan range: m/z 50-1,000

Sheath liquid: HMT Sheath Liquid (p/n : H3301-1020)

<u>Anionic Metabolites (Anion Mode)</u>:

Run buffer: Anion Buffer Solution (p/n : H3302-1021)

Rinse buffer: Anion Buffer Solution (p/n : H3302-1021)

Sample injection: Pressure injection 50 mbar, 25 sec

CE voltage: Positive, 30 kV

MS ionization: ESI Negative

MS capillary voltage: 3,500 V

MS scan range: m/z 50-1,000

Sheath liquid: HMT Sheath Liquid (p/n : H3301-1020)

Data Processing and Analysis

Data processing and analysis were done by Human Metabolome Technologies as described before^21–23^. Briefly, MasterHands ver. 2.17.1.11 software was used to extract peaks detected in CE-TOFMS analysis in order to obtain m/z, migration time (MT), and peak area. Relative peak area was calculated as:

> Relative Peak Area = Metabolite Peak Area / (Internal Standard Peak Area × Sample Amount)
>
> Signal-noise ratio (S/N) = 3 was set as a threshold for peak detection limit.

Human Metabolome Technologies target library was used to identify putative metabolites and their isotopic ions on the basis of m/z and MT with tolerance ±0.5 min in MT.

Absolute quantification was performed in total amount of each detected metabolite. All the metabolite concentrations were calculated by normalizing the peak area of each metabolite with respect to the area of the internal standard and by using standard curves, which were obtained by single-point (100 μM) calibrations.

### pH Measurements

Five milliliters of the medium were centrifuged at 500 Xg for 5 minutes to remove suspended cells and the pH was measured using a pH meter (Mettler Toledo, Colombus, OH)

### Mitotracker Green and TMRE Assays

MitoTracker™ Green FM (Thermofisher # M7514) or Tetramethylrhodamine ethyl ester perchlorate (TMRE, Thermofisher # T669) was added directly to the medium to a final concentration of 200 nM or 50 nM respectively. After 20 minutes incubation at 37°C, the medium was removed and the cells were washed twice in 1X PBS and harvested. Fluorescent intensity was measured in at least 20,000 cells as suggested by the manufacturer. Data are presented as a geomean ± 95% confidence interval. The comparison between the fluorescence intensity in cells with or without doxycycline was done using the two-tailed T-Test.

### Seahorse Assay

#### Seahorse measurement of mitochondrial function and glycolytic flux

Mitochondrial function and glycolytic flux were determined using Seahorse XFe 96 extracellular flux analyzer (Seahorse, Agilent) as oxygen consumption rate (OCR) and extracellular acidification rate (ECAR) using “Mito-stress test” or “Glycolysis stress test” methods, respectively. Cells harboring inducible N-Myc overexpression vector were treated +/− DOX for 2, 3, 4 or 5 days. Next, cells were plated onto 96-well plate using several strategies to verify experimental outcome. All plating approaches yielded consistently very similar results. In particular, cells were plated at two different concentrations (4×10^4^ or 2.5×10^4^ cells/well) in multiple wells (i) allowing to attach for 6 h or (ii) over-night to fibronectin-coated wells or (iii) were attached readily after re-plating using Cell Tak cell adhesive (Corning). FBS-free Seahorse XF RPMI medium (without Phenol Red, Agilent) was supplemented with 2 mM glutamine, 1 mM sodium pyruvate and 2 g/L glucose (for “Mito-stress test”) or with 2 mM glutamine (for “Glycolysis stress test”). Measurements were performed at basal state and following serial injections of: oligomycin (1 μM), FCCP (0.5 μM) and rotenone/antymycin A (0.5 μM/1 μM) for “Mito-stress test” or of glucose (10 mM), oligomycin (1 μM) and 2-deoxy-glucose (2-DG, 20 mM) for “Glycolysis stress test”.

#### TEM Analysis of Mitochondria

The samples were fixed with 2.5% glutaraldehyde, 2% paraformaldehyde in 0.1M sodium cacodylate buffer pH 7.4 and post fixed in 2% osmium tetroxide in 0.1M cacodylate buffer with 0.3% potassium ferrocyanide for 1.5 hours. After rinsing in the same buffer, the tissue was dehydrated through a series of graded ethanol to propylene oxide, infiltrated and embedded in epoxy resin and polymerized at 70°C overnight. Semithin sections (0.5micron) were stained with toluidine blue for light microscope examination. Ultrathin sections (80nm) were cut and imaged using an FEI Tecnai 200Kv FEG Electron Microscope with an ATM XR41 Digital Camera. To score mitochondrial damage, 50 images from every sample were selected in which the nucleus and at least 5 mitochondria were visible at 5K magnification. Images of this profiles were taken at 29K with some portion of the nucleus visible, printed and all mitochondria profiles were scored blindly. Mitochondria were scored 0= normal (aligned cristi and normal outer membrane), 1=abnormal (swollen and disorganized cristi or disrupted outer membrane) or 2= Very abnormal (both swollen and disorganized cristi and severe outer membrane disruption).

#### Gamma H2AX Immunostaining

Cells were harvested and incubated on poly-L-Lysin-treated (Sigma, Cat. # 8920) slides for 30 minutes at 37°C. The cells were fixed in 4% paraformaldehyde in PBS at 4°C overnight, blocked by normal donkey serum for 1 hour at room temperature and incubated with mouse anti Phospho Ser139-histone H2AX antibody (Millipore, Cat. # 5-636) 1:10,000 overnight at 4 °C. The slides were then washed 3 times in PBS and incubated with donkey anti-mouse Cy3 IgG antibodies (Millipore, Cat. # AP192C) 1:500 for 1 hour at room temperature. The slides were washed 3 times in PBS, counter-stained with DAPI for 10 minutes, washed 3 times in PBS and mounted with Prolong™ Gold Antifade Mountant (Thermo Fischer, Cat. # P36930).

#### Measuring ROS

Reactive Oxygen Species (ROS) was assayed using DCFDA cellular ROS detection assay kit (Abcam # ab113851) or CellRox Green (ThermoFischer # C10444) following manufacturers’ protocol. For DCFDA cellular ROS detection assay kit, cells were plated overnight in 96 well plates (25,000 cells per well) in their complete medium with or without doxycycline. The fluorescence excitation/ emission 485/535 was measured using the microplate reader SpectraMax-M5 (Molecular Devices, Sunnyvale, California). For CellRox Green, cells grow in 10-cm plates. CellRox Green was added to the medium 1:500 and incubated for 30 minutes then cells were harvested and analyzed using flowcytometry.

#### Measuring cellular Glutathione

The cells were plated in 96 well plates overnight at a density of 6,000 cells per well in their complete medium with or without doxycycline. Glutathione concentration per well was measured using GSH-Glo Glutathione Assay Kit (Promega # V6911) according to the manufacturer’s protocol. The luminescence was measured using PHERAstar FSX (BMG Labtech, San Diego, California)

#### Drug screening

Cell screening was done on MYCN-induced and un-induced SKNMM^MYCN^ cells as previously described^6^ with the following modifications: to determine optimal cell plating density, SKNMM^MYCN^ cells was plated on a flat-bottomed, white 96-well plate (Corning Cat#3917), at eight different cell densities ranging from 0 cells/well to 10,000 cells/well; 12 wells per cell intensity per plate. Twenty-four hours after plating, medium was changed with or without doxycycline. Medium was changed after that with or without doxycycline every 2 or 4 days. One plate from every testing condition was read every day for 11 days using CellTiter-Glo^®^ (‘CG’, Promega Cat#G7573) on PHERAstar FSX (BMG Labtech, San Diego, California). Cell density of 1,250 cells per well and medium change once every 4 days were determined to have good signal to noise separation while maintaining logarithmic growth in both induced and un-induced cells. We tested different DMSO concentrations for the selected cell density and confirmed minimal cell death with up to 0.197% DMSO. Positive control compound selection was performed with eight positive control compound candidates (doxorubicin HLC, staurosporine, etoposide, SN-38, bortezomib, cyclohexamide, panobinostat and TAKA123) arrayed in single point concentration and 1:3 dilution series. Based on successful cell killing Staurosporine was chosen as positive control for cell screenings. Assay validation in 384 well plates were performed with staurosporine.

#### Dose Response curve experiment

A schematic representation of the dose response curve experiment is illustrated below. Cells were plated in 96 well plates (Corning Cat#3917) at 1,250 cells/100μL; twelve cell plates per compound plate. Approximately 24 hours following plating, medium was changed with fresh medium with or without doxycycline; 6 cell plates per a compound plate each. Two days after MYCN induction, medium was changed again with fresh medium with or without doxycycline and cells were drugged with the compound plates and a positive control plates using a Biomek FX (Beckman Coulter) liquid handler equipped with a pin tool. The pin tool transferred 112 nL compound stock, resulting in 890-fold compound dilution. After 3 days of drugging, half of the plates were read. On day 4 post drugging, medium was changed with fresh medium again with or without doxycycline and fresh drugs were added to the medium using the Biomek FX. After 4 days of the second drugging, the other half of the plates were read. The compound plate contained compounds dissolved in DMSO arrayed in a 1:3-fold dilution series in columns 1-10 with 8 compounds per plate (target concentration 10mM or 1mM). The positive control plate was empty from columns 1-10; in columns 11-12, it contained DMSO and the positive control. Two biological replicates were performed for each 8-days experiment, with three technical replicates per biological replicate.

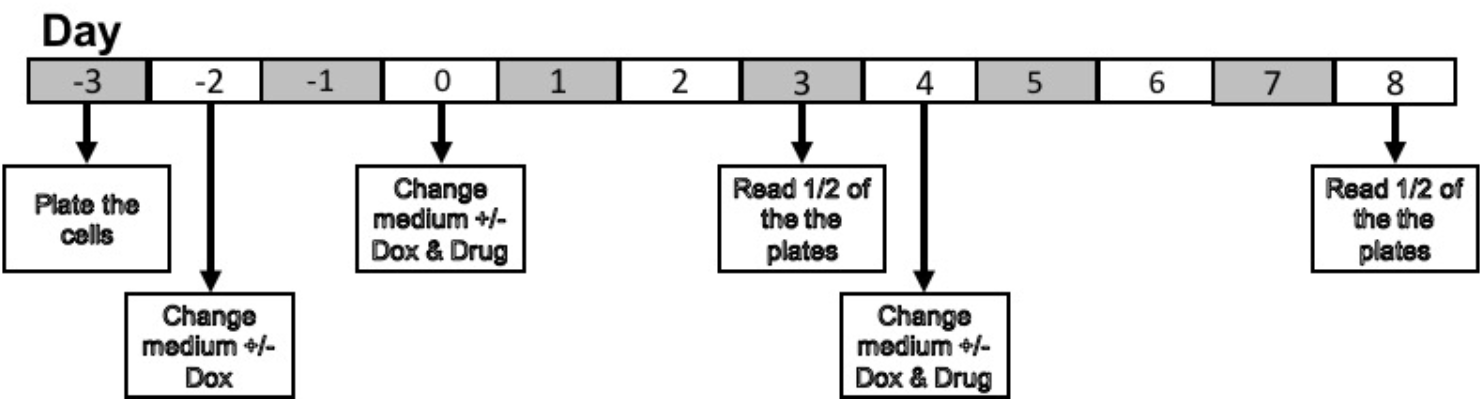

#### Cell screening analysis

Normalization, fitting, and visualization of dose-response experiments were performed using custom code written in the R programing language (version 3.3.2, R Core Team (2016). R: A language and environment for statistical computing. R Foundation for Statistical Computing, Vienna, Austria. URL https://www.R-project.org/.).

Dose response curve data analysis for cytotoxic compounds was done as previously described before^6^. Raw CellTiter-Glo (CTG) luminescence signals (RLU) were first log2 transformed, then normalized to the mean of the positive and negative controls on each plate.

In order to detect drugs that reduced MYCN-induced cell death, traditional dose-response analysis was not appropriate because protective drugs would often protect at low concentrations, but eventually induce cytotoxicity at high concentration. To detect a protective effect, we needed to quantify the increase in CTG signal at low concentration before any decrease in signal at high concentration. To do so, the CTG RLUs from all wells were first log2 transformed for each plate, the value for each well was then normalized by subtracting out the mean log2 RLU for negative control wells. The normalized activity of each compound was then fit as a function of the log10 drug concentration using a smooth spline (smooth.spline function in R with default parameters). For each fit, the area under the curve (AUC) for the portion of the spline with RLU above the RLU of the lowest drug concentration tested was calculated. Higher values for this AUC metric indicate greater protection against loss of cell viability as measured by the CTG assay.

#### CUX-2 expression in SKNMM^MYCN^ cells

SKNMM^MYCN^ cells were transduced by a lentiviral vector (pLenti-C-mGFP-P2A-Puro) from Origene expressing either CUX-2 (Cat# RC222063L4V) or GFP control construct (Cat# PS100093V). GFP-positive cells were sorted after 48 hours and maintained in a selective media containing 1.5-ug puromycin per ml.

